# A critical developmental interval of coupling axon branching to synaptic degradation during neural circuit formation

**DOI:** 10.1101/2022.04.06.486606

**Authors:** Suchetana B. Dutta, Gerit Arne Linneweber, Maheva Andriatsilavo, Peter Robin Hiesinger, Bassem A Hassan

## Abstract

The emergence of neuronal wiring specificity requires stabilization of dynamic axonal branches at sites of selective synapse formation. Models that explain how axonal branching is coupled to synaptogenesis postulate molecular regulators acting in a spatiotemporally restricted fashion. We report that Epidermal Growth Factor Receptor (EGFR) activity is required in presynaptic axonal branches during two distinct temporal intervals to regulate circuit wiring in the developing *Drosophila* visual system. EGFR is required early to regulate primary axonal branching and independently again later to prevent autophagic degradation of the synaptic active zone protein Bruchpilot (Brp). The protection of synaptic material during this later interval of wiring ensures the stabilization of terminal branches, circuit connectivity and appropriate visual behavior. Phenotypes of EGFR inactivation were rescued by increasing Brp levels or downregulating autophagic genes. We identify a temporally restricted molecular mechanism required for coupling axonal branching and synaptic stabilization that contributes to the emergence of neuronal wiring specificity.

## INTRODUCTION

Stereotypy and robustness of neuronal circuit wiring is thought to be critical for normal brain function, and malformations of neuronal circuits are associated with a spectrum of neurodevelopmental and neuropsychiatric diseases like Autism Spectrum Disorder and Schizophrenia (Powchik *et al*., 1998; Doll and Broadie, 2014; Moyer, Shelton and Sweet, 2015). Whereas the outcome of axonal branching and synapse formation is spatially and temporally highly stereotyped, the developmental process itself relies heavily on probabilistic local events like filopodial growth and retraction, local protein recycling and degradation, self-avoidance of exploratory filopodia through non-deterministic expression of cell adhesion molecules, cytoskeletal polymerization and depolymerization, mitochondrial fusion and fission, and stochastic synaptic seeding (Chen *et al*., 2006; Zschätzsch *et al*., 2014; Winkle *et al*., 2016; Simmons *et al*., 2017; Constance *et al*., 2018; Lewis *et al*., 2018; M. Neset Özel *et al*., 2019; Urwyler *et al*., 2019). This raises the question of how such events are orchestrated in space and time to reproduce stereotyped circuit wiring diagrams. The formation of functional neuronal circuits relies on neurons branching their axons during development in an orderly spatial and temporal manner to connect with a specific set of post-synaptic neurons (Rico *et al*., 2004; Colón-Ramos, 2009; Kalil and Dent, 2014; Batool *et al*., 2019; Agi, Kulkarni and Hiesinger, 2020) but how individual axons and their branches locally pattern their connectivity to their post-synaptic targets is not well understood. Evidence from various systems shows that during late stages of neuronal circuit formation, after an initial phase of exploratory axonal branching, interdependence between synapse formation and branching dynamics plays a key role in the selection of future synaptic partners (Meyer and Smith, 2006; Ruthazer, Li and Cline, 2006; Chia *et al*., 2014; Kalil and Dent, 2014; Xu and Quinn, 2016; Constance *et al*., 2018). This ‘synaptotropic’ iterative interaction between branching and synapse formation ensures reproducibility of circuit wiring patterns (Rico *et al*., 2004; Chia *et al*., 2014) and might itself act as a limiting factor during development to prevent excessive branch growth resulting in a stable adult pattern (Niell, 2006). These findings necessitate the existence of local molecular mechanisms that act in a temporally specific manner to couple axonal branching dynamics to synapse formation. The identity and mode of action of such temporal molecular coupling events are poorly understood.

The formation of a stable synaptic contact is a function of the equilibrium between synaptic seeding and synaptic degradation as shown by developmental pruning of up to 40%-50% of synapses (Bourgeois and Rakic, 1993). Synapse elimination is an important cellular phenomenon which fine tunes neural circuitry (Hua and Smith, 2004) especially during the widely studied postnatal experience dependent plasticity known the as “critical period” (Cisneros-Franco *et al*., 2020). Interestingly, there is evidence for what has bene referred to as “precritical period” plasticity in the visual cortex (Feller and Scanziani, 2005), suggesting that genetically encoded developmental events may define critical developmental intervals of synaptic consolidation and elimination prior to the onset of experience dependent plasticity. The timing, role and molecular regulation of such developmental events are unknown.

Autophagy is a key cellular homeostasis mechanism, the dysregulation of which is thought to contribute to several neurodegenerative and neurodevelopmental diseases (Poultney *et al*., 2013; Huber *et al*., 2015; Kim *et al*., 2016; Menzies *et al*., 2017; Vijayan and Verstreken, 2017; Stavoe and Holzbaur, 2019). Recent work has established that autophagy also plays a role in brain development. Loss of autophagy results in morphological and functional presynaptic organization defects in wide-range of systems, including the mouse cochlear ribbon synapse (Xiong *et al*., 2020) and the *Drosophila* mushroom body and photoreceptor neurons, for example (Bhukel *et al*., 2019; Fleming and Rubinsztein, 2020; Kiral *et al*., 2020). At the *Drosophila* neuromuscular junction, disruption of autophagy reduces its size whereas induction of autophagy increases synaptic boutons and neuronal branches (Shen and Ganetzky, 2009). Autophagy deficiency also causes dendritic spine pruning defects and Autism-like social behaviors in a mouse model (Tang *et al*., 2014). Pre-synaptic sites are zones of autophagosome biogenesis that have been linked to synaptic plasticity (Azarnia Tehran, Kuijpers and Haucke, 2018; Wang *et al*., 2019). Local autophagy may also play a role in positioning axonal branches (Adnan *et al*., 2020). How axonal branch growth and refinement are molecularly coupled to synapse formation and pruning in space and time during neuronal circuit wiring is not well understood.

We have previously identified a role for local activation and recycling of the Epidermal Growth Factor Receptor (EGFR) in regulating dynamics and final patterning of axonal branches (Zschätzsch *et al*., 2014) in higher order object response neurons (Linneweber *et al*., 2020) in the fly visual system. Dorsal Cluster Neurons (DCN) are a bilateral cluster of 22-68 commissural interneurons in the *Drosophila* brain with their cell bodies in the dorso-lateral portion of the central brain and axons crossing the central brain to project onto contralateral optic lobes. A subset of DCNs, referred to as M-DCNs, project their axons to a distal visual neuropil called the Medulla where they form a stereotypical fan shaped branching pattern (Srahna *et al*., 2006; Langen *et al*., 2013). M-DCN wiring patterns determine visual object orientation behavior in flies. Local asymmetric localization and recycling of EGFR in M-DCN axonal filopodia during development has been linked with actin polymerization and filopodial dynamics underlying the patterning of presynaptic M-DCN branches. This offers a convenient model system and a molecular entry point to dissect the mechanisms that link axonal branching to synapse formation during neuronal circuit development resulting in specific wiring patterns underlying behavior.

We investigated whether and how M-DCN axonal branching is molecularly coupled to synapse formation and what the consequences to circuit wiring and behavior might be if this link is disrupted. We found that EGFR activity is required at two distinct temporal intervals: an early actin-dependent interval to establish primary axonal branches where presynaptic material accumulates, and a later interval to prevent autophagic degradation of the presynaptic active zone scaffold protein Bruchpilot (Brp) and allow stabilization of the synaptic contacts. Therefore, a temporal sequence of local molecular interactions coordinated by EGFR signaling ensures the coupling between progressive axonal branch refinement and stabilization of the presynaptic active zone leading to the emergence of axon-specific connectivity.

## RESULTS

### Spatio-temporal correlation of the molecular mechanisms of branching and synapse formation during development

To investigate the relationship between terminal axon branching and synapse formation at high spatio-temporal resolution, we used the medulla innervating Dorsal Cluster Neurons (M-DCNs) as a model. M-DCN axons form stable, terminal primary (arrowhead) and secondary (arrow) presynaptic branches in several posterior medulla layers with their dendrites projecting to the ipsilateral lobula (Zschätzsch *et al*., 2014) (Fig.1A,A’). Each M-DCN axon forms an average of 4.5 primary branches/axon and an average of 0.25 secondary branches/primary branch (Fig.S1G). These terminal branches contain the presynaptic sites of M-DCNs as marked by several presynaptic and active zone markers such as Syt, Syd1 and Brp (Fig.1A,A’-C,C’). We have previously shown that the number of M-DCN axonal presynaptic branches is regulated by local EGFR activity (Zschätzsch *et al*., 2014) between 48 and 72 hours of pupal development (P48-P72) at 25 degrees.

**Fig.1.**
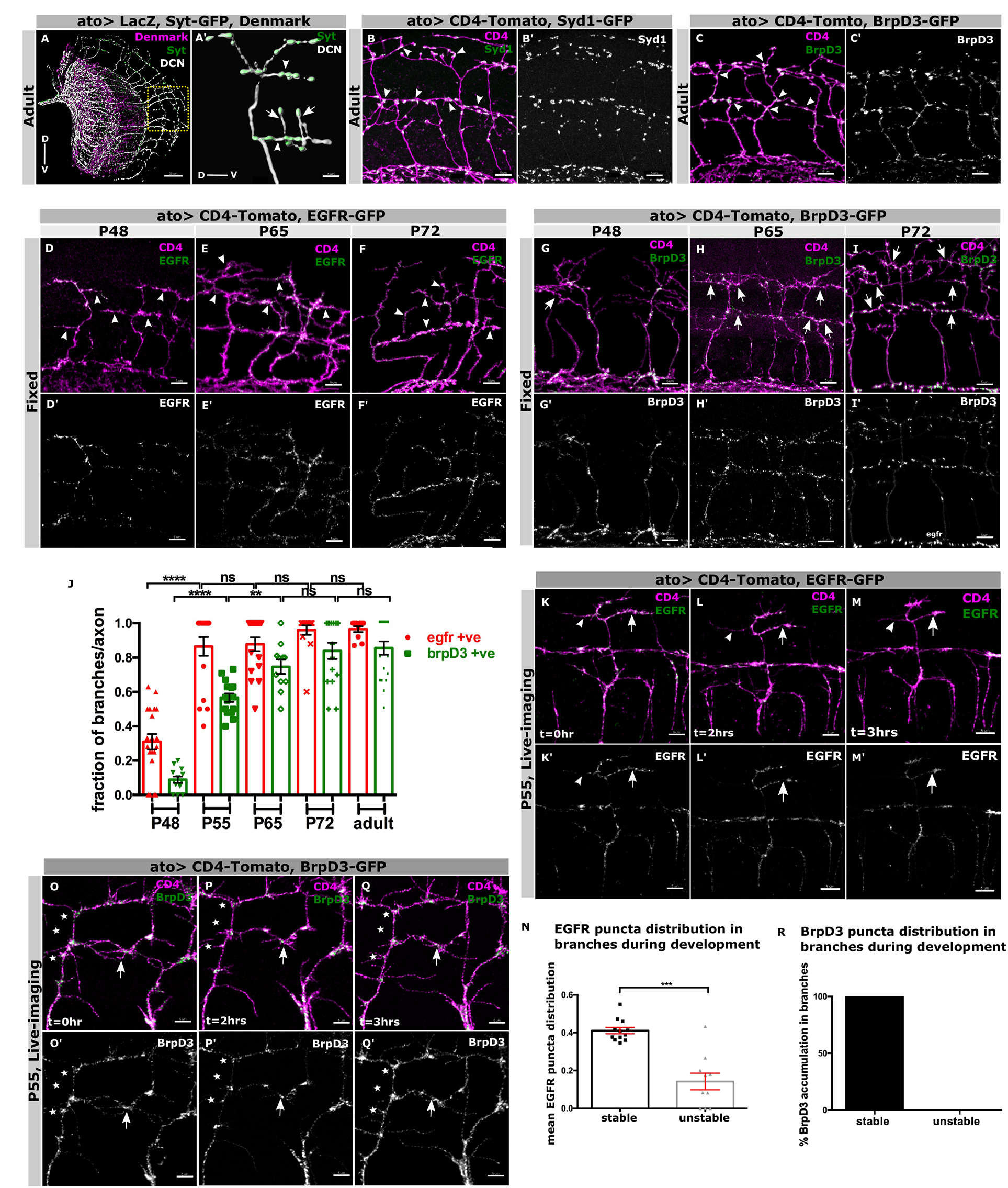
Spatio-temporal correlation of the molecular mechanisms of branching and synapse formation during development: **(A-A’)** Adult stereotypic projection pattern of DCNs driven by *atoGal4-14a* labelled with LacZ (white), dendritic arborizations marked by Denmark (magenta) and pre-synaptic sites marked by Syt-GFP (green). *atoGal4-14a* is used in all the following experiments unless otherwise stated. Yellow box represents region of interest where DCN axons forms ladder-like branches as shown in **(A’)** higher resolution with primary (asterisks) and secondary branches (arrowheads) harboring Syt-GFP (green) puncta. **(B/B’-C/C’)** Adult DCN branches (magenta) are associated with clusters of early synaptic seeding factor; marked with Syd1 GFP (green) **(B/B’)** and late Active zone (AZ) protein Brp, marked with BrpD3 GFP (green) **(C/C’). (D-I)** Temporal order of recruitment of EGFR GFP (green) and BrpD3 GFP (green) in the DCN branches (magenta) during development; P48 **(D/D’**,**G/G’)**, P65 **(E/E’**,**H/H’)** and P72 **(F/F’**,**I/I’)** showing EGFR enters the branches before Brp. **(J)** Quantification showing the fraction of branches recruiting EGFR GFP (red datapoints) and BrpD3 GFP (green datapoints) during development. N=20 axons at P48 (EGFR), N=17 axons for at P48 (BrpD3), N=17 axons at P55 (EGFR), N=17 axons at P55(BrpD3), N=17 axons at P65 (EGFR), N=14 axons at P65 (BrpD3), N=15 axons at P72 (EGFR), N=16 axons at P72 (BrpD3), N=10 axons in adult (EGFR) and N=20 axons in adult (BrpD3). Mann Whitney test; \*\*\*\**p* < 0.0001, ^**ns**^*p*=0.8124, ^**ns**^*p*=0.7934, ^**ns**^*p*=0.4932. Ex-vivo imaging of EGFR GFP (**K/K’-M/M’)** and BrpD3 GFP **(O/O’-Q/Q’)** in the DCN branches (magenta) during development shows EGFR GFP (green) asymmetrically localizes in the DCN branches; stable branches with higher proportion of EGFR punctas (arrow) compared to unstable branches (arrowhead)**(K/K’-M/M’)**. Whereas late AZs marked by BrpD3 GFP (green) accumulates only in stable branches (arrow), while being excluded from unstable branches (arrowhead) **(O/O’-Q/Q’). (N)** Quantification of mean EGFR GFP puncta normalized to branch length in stable (black bar) vs unstable branches (gray bar) in ex-vivo cultures during development. N=12 for stable branches and N=10 for unstable branches, Mann Whitney test, \*\*\**p* =0.0003. **(R)** Quantification of percentage of BrpD3 GFP puncta accumulation in stable (black bar) vs unstable branches during development. N=10 for stable branches and N=15 for unstable branches. Error bars denote mean±SEM, scalebar represents 5μm except (A) which is 20μm.

To investigate the spatio-temporal pattern of synapse formation in the context of axonal branching, we quantified the order of appearance of EGFR (EGFR-GFP), Syd1 (Syd1-GFP) and Brp (BrpD3-GFP) in M-DCN branches between P48 and P72 using well-established reporters of the localization of the endogenous proteins that show no known overexpression phenotypes (Fouquet *et al*., 2009; Owald *et al*., 2010; Zschätzsch *et al*., 2014). At P48, ∼60% of the branches contain Syd1-GFP (Fig.S1A, A’,D), ∼30% contain EGFR-GFP, but less than 10% contain BrpD3 (Fig.1D,D’,G,G’,J). Between P55 and P65 almost all branches contain Syd1-GFP (Fig.S1B,B’,D), ∼90% contain EGFR-GFP and BrpD3 increases from ∼50% at P55 to ∼75% at P65 (Fig.1E,E’,H,H’,D). By P72, all three proteins have reached their stable adult levels (Fig.1F,F’,I,I’,J; Fig.S1C-D). In conclusion, Syd1 enters DCN axons much earlier than EGFR prior to branch formation at around P40 (Fig.S1E,E’-F,F’), followed by Brp entry. To reveal the dynamics of this process, we live imaged and quantified the trafficking of discrete clusters (henceforth “puncta”) of BrpD3-GFP, Syd-1-GFP and EGFR GFP in *ex vivo* cultures at P55 when all three proteins are present in branches *in vivo* and EGFR is known to regulate branch growth and pruning (Zschätzsch *et al*., 2014). We defined any branch as stable that was present during the entire imaging session (∼8hours), and as unstable any branch that retracted without re-growing during the same time. Whereas Syd1 entered all branches regardless of stability (FigS1H,H’-J,J’,K, Supplemental Video S1), EGFR accumulated preferentially, but not exclusively, in stable branches (arrow) compared to unstable branches (arrowhead) (Fig.1K,K’-M,M’,N, Supplemental Video S1). Brp exclusively accumulated in stable branches (arrow), despite being present throughout the entire axon shaft (asterisks). None of the branches which failed to accumulate BrpD3 puncta during development stabilized, suggesting active zone maturation is a prerequisite for branch stabilization (Fig.1O,O’-Q,Q’, R, Supplemental Video S1). Taken together, our spatial and temporal analyses of *in vivo* and live imaging data suggest that almost all branches are synapse competent (contain Syd1 early), but only the fraction that accumulates both EGFR and Brp is stabilized to contain future synaptic active zones.

### Synapse formation is required for branch patterning

To investigate the molecular interdependence of axon branching and synapse formation, we inactivated EGFR, Syd1 or Brp and examined the development of the branches *in vivo* starting at P48. In controls, M-DCN axon branch numbers increased gradually over time between P48 and P65 and then began to decline, reaching near adult branch numbers at P72 (Fig.2A-A5, C). This gradual branch refinement between P65 and P72, follows Brp entry into stable branches (Fig.1J, R), suggesting a link between final branch pruning and stabilization under wild type conditions. As previously shown, we found that inactivating EGFR using a well-established dominant negative transgene (EGFR-DN) (Buff *et al*., 1998) resulted in an increase in the total number of branches in adult (Fig.2C). Surprisingly however, loss of EGFR function changed the temporal pattern of branch pruning from a gradual increase followed by a gradual decrease, into a bi-phasic growth and pruning (Fig.2B-B5,C) mode affecting both primary and secondary axon branches (Fig.2D-E). We had previously shown that loss of EGFR function causes primary branch pruning defects between P48 and P55 through an actin-dependent mechanism (Zschätzsch *et al*., 2014); whereas our data suggests a surge in secondary branch outgrowth between P65-P72 as the possible reason behind second branching peak during late development (Fig.S2G, Supplemental Video S2.1). We therefore asked whether both the primary and secondary branching phenotypes of EGFR inactivation at two distinct phases can be explained by changes to actin dynamics. We found that RNAi knockdown of several cytoskeletal regulators resulted in an increase in in primary branches, but not in secondary branches (Fig.S2A-F). Together, these data suggest that EGFR is required at two distinct temporal phases for DCN branch dynamics. The second phase, which starts at P65, appears to be mechanistically distinct from the first, and coincides with Brp recruitment into stable branches. We therefore asked whether loss of synaptic proteins influences axonal branching specifically during the second phase of branch outgrowth. We knocked down Syd-1 and Brp specifically in DCNs (using ato-Gal4) throughout development using Syd1 RNAi and Brp RNAi (line B3,C8;Wagh et al., 2006). Like EGFR inactivation, knockdown of both Syd1 and especially Brp, resulted in an increase in the numbers of primary and secondary branches in adults (Fig.2C,F-K). However, the branch pruning defects caused by loss of Syd1 and Brp only overlapped with the second phase of EGFR requirement between P65 and P72 (Fig.2C) with similar increased secondary branching during late development starting P65 in Brp KD (Fig.S2G, Supplemental Video S2.1). Finally, downregulation of Brp resulted in continued exploratory growth and retraction even in adult brains (Supplemental Video S2.2). These observations suggest a temporally restricted link between EGFR activity, branching dynamics and synapse formation during a late developmental interval of neural circuit wiring.

**Fig.2.**
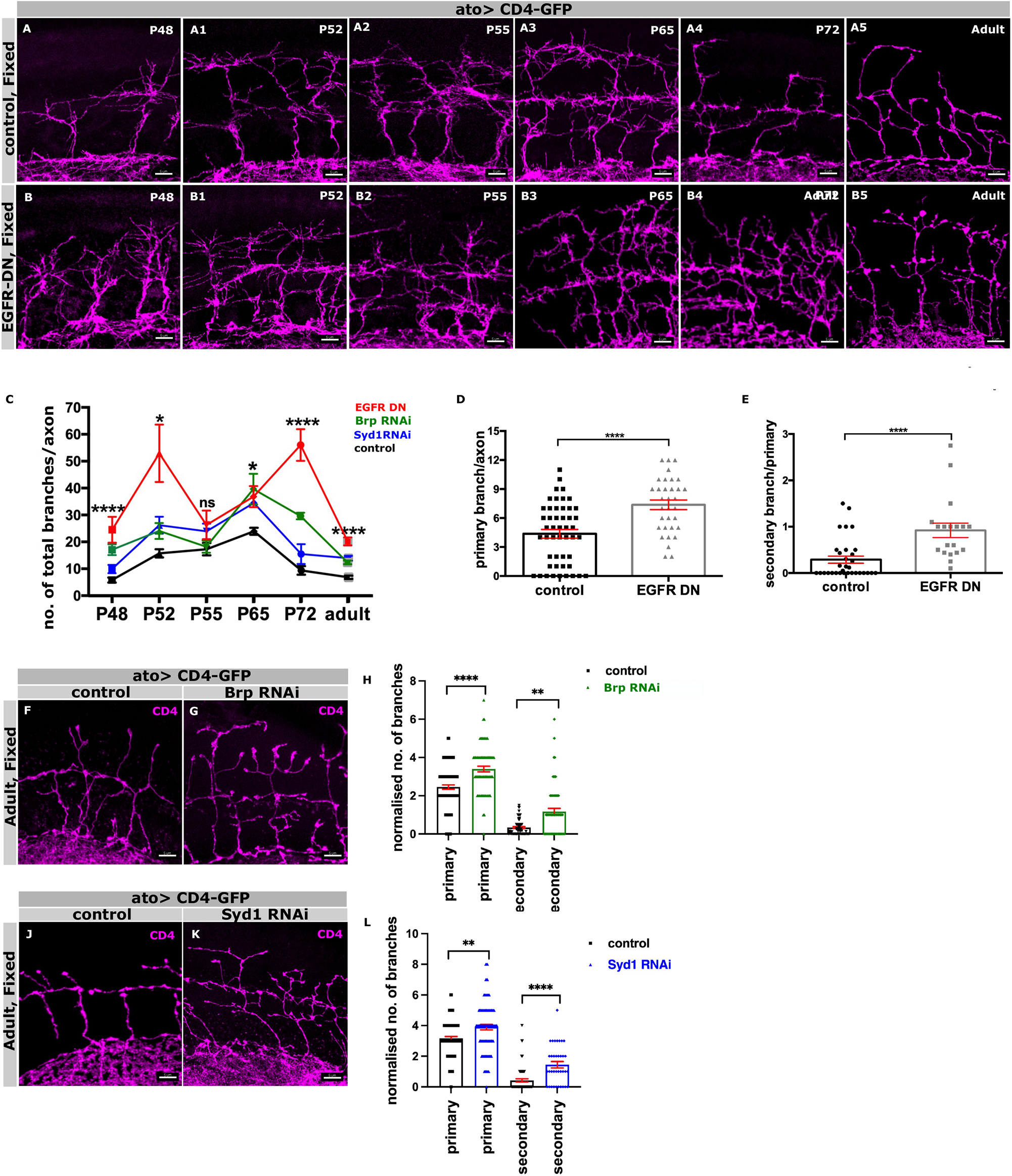
Synapse formation is required for terminal branch patterning: **(A-A5)** DCN axonal branch progression (magenta) during development in control brains reveals an increase in branching complexity from early to mid-pupal stage; P48 **(A)**, P52 **(A1**), P55 **(A2)**, P65 **(A3)**. Branch refinement occurs from mid to late-pupal development; P65 **(A3)**, P72 **(A4)**, adult **(A5). (B-B5)** Surprisingly, EGFR DN expressing DCNs show dys-regulation of branch refinement process with branching peaks at two phases of development: first from P48 to P55 **(B-B2)** and later from P65 to adult **(B3-B5). (C)** Quantification showing overall branch refinement progression (mean of the total number of branches per axon) during development in EGFR DN (red), Brp RNAi (green), Syd1 RNAi (blue) compared to control (black). N=8 lobes for control, N=10 lobes for EGFR DN, N=7 lobes for Syd1 RNAi and N=6 lobes for Brp RNAi. Kruskal–Wallis and Dunn’s as post-hoc test; \*\**p* =0.0015(P48), \*\**p* =0.0033(P52), ^***ns***^*p* =0.2135(P55), \*\**p* =0.0024(P65), \*\*\**p* =0.0002(P72) and \*\*\**p* =0.0004(adult). **(D)** Quantification showing more normalized primary branch number in EGFR DN (gray) expressing DCNs compared to genetic control (black) in adults. N=49 axons for control, N=33 axons for EGFR DN; T-test, \*\*\*\**p* < 0.0001. **(E)** Quantification showing more normalized secondary branch number in EGFR DN (gray) expressing DCNs compared to control (black) in adults. N=32 primary branches for control, N=19 primary branches for EGFR DN; T-test, \*\*\*\**p* < 0.0001. Error bars denote mean±SEM. **(F-G, J-K)** Adult axonal branch pattern (magenta) upon knocking down Brp **(F-G)** and Syd1 **(J-K)** specifically in DCNs compared to their genetic controls. **(H-I)** Adult quantification showing normalized primary **(H**) and secondary **(I)** branch number increases in Brp B3,C8 RNAi (green) compared to control (black). N=49 axons for control, N=38 axons for Brp B3,C8-RNAi for primary branch quantification **(H)**, N=54 branches in control, N=60 branches for secondary branch quantification in Brp B3,C8 RNAi **(I)**. Mann Whitney test, \*\*\*\**p* < 0.0001, \*\**p* = 0.0011. **(L-M)** Adult quantification showing similar increase in normalized primary **(L)** and secondary **(M)** branch number of adult DCNs in Syd1 RNAi (blue) compared to control (black). N=92 axons for control, N=90 axons for Syd1-RNAi **(L)**, N=66 branches in control, N=36 branches in Syd1-RNAi **(M)**. Mann Whitney test, \*\**p* = 0.0011, \*\*\*\**p* < 0.0001. Error bars denote mean±SEM, scalebar represents 5μm.

### EGFR is required for Brp stabilization and synaptic connectivity

To cross-regulation between EGFR activity and synapse formation, we asked whether and how Syd1, EGFR, and Brp interact genetically to establish the pattern of M-DCN connectivity. Syd1 is known to recruit Brp (Spinner, Walla and Herman, 2018) to future presynaptic sites (Owald *et al*., 2010). However, knockdown of Syd1 had no effect on the distribution or levels of EGFR, neither did knockdown of Brp (Fig.S3A,A’-C,C’,D). Conversely, inactivation of EGFR did not affect Syd1 levels or distribution (Fig.3A,A’-B,B’, E). This suggests that EGFR and Syd1 act in parallel during synapse formation. In contrast, loss of EGFR activity resulted in a significant decrease of BrpD3-GFP puncta – which colocalize with endogenous Brp (Fig.S3E,E’-F,F’) – in terminal branches from 0.22 puncta per µm to 0.1per µm. This corresponds to a 2.5-fold decrease in the total number of BrpD3 puncta from ∼15 per axon to ∼6.5 per axon and was accompanied by a redistribution from numerous small puncta at presynaptic sites into a few large BrpD3 aggregates (arrow) (Fig.3C,C’-D,D’,F,G), an indicator of loss of endogenous active zones in DCNs (Kiral *et al*., 2021).

**Fig.3.**
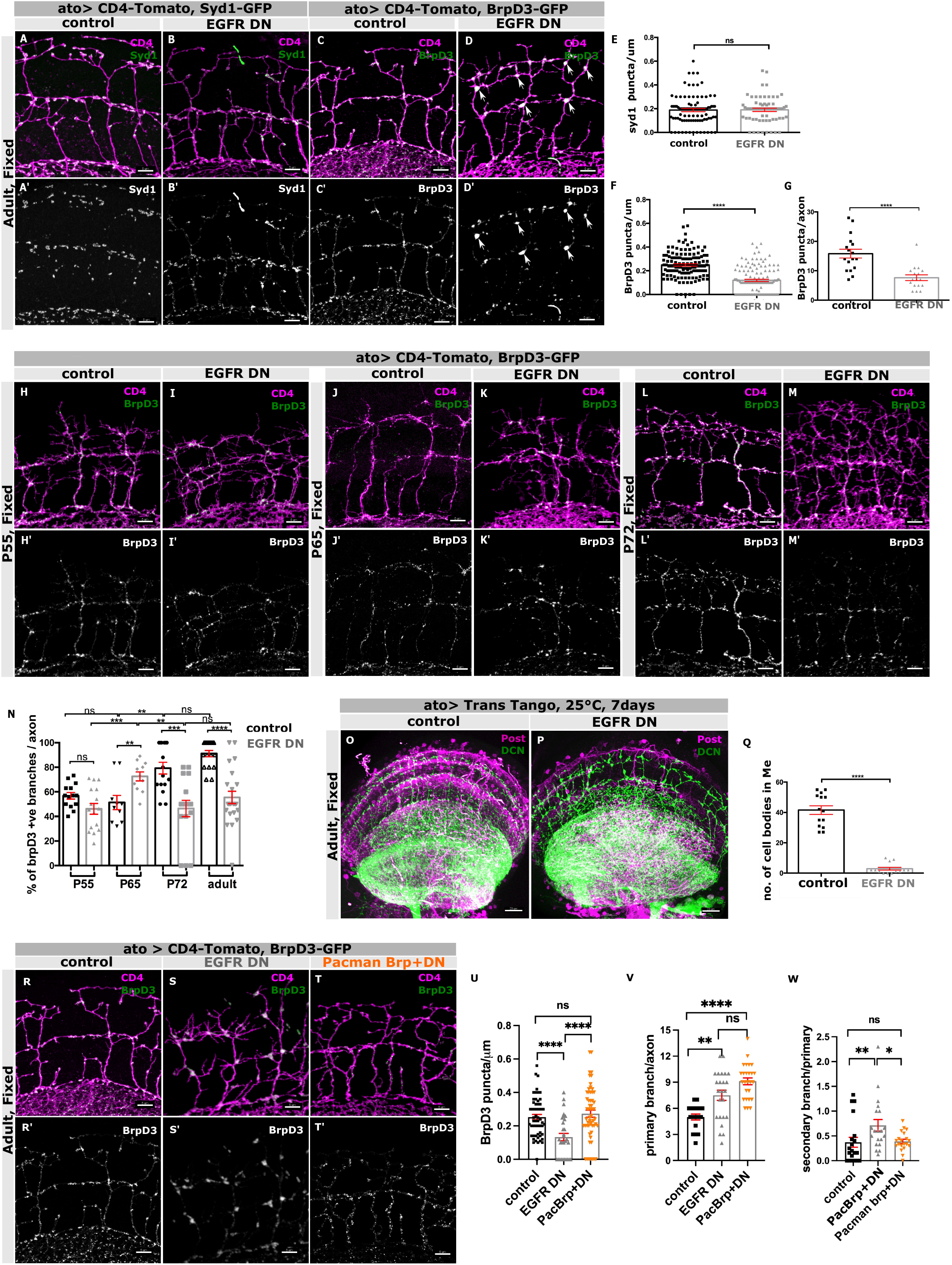
EGFR is required for Brp stabilization in terminal branches: **(A/A’-D/D’)** Adult DCN axon branches contain early seeding factor Syd1-GFP which is unaffected by EGFR-DN **(A/A’-B/B’)** while late AZ marker BrpD3-GFP reduces drastically in the branches where they accumulate in larger volumes (white arrows) in EGFR DN **(C/C’-D/D’**). **(E)** Quantification showing unaffected distribution of Syd1-GFP puncta normalized to axon branch length in control (black) vs EGFR DN (gray) in adults. N=93 branches for control and N=66 branches for EGFR DN. T-test, *p* =0.6688. **(F)** Quantification showing the reduced distribution of BrpD3-GFP puncta normalized to axon branch length in control (black) vs EGFR^DN^ (gray) in adults. N=124 branches for control and N=124 branches for EGFR DN, T-test, \*\*\*\**p* < 0.0001. **(G)** Adult quantification of the total number of BrpD3-GFP puncta per axon in control (black) vs EGFR DN (gray). N=17 axons for control and N=17 for EGFR DN. Mann Whitney test; \*\*\*\**p* < 0.0001. **(H/H’-M/M’)** Recruitment of BrpD3-GFP puncta (green) in developing DCN branches labeled by CD4-tomato (magenta) in fixed samples during different stages of pupal development in control vs EGFR DN; P55 **(H/H’-I/I’)**, P65 **(J/J’-K/K’)** and P72 **(L/L’-M/M’). (N)** Quantification showing percentage of BrpD3-GFP positive DCN branches per axon during development in control (black) vs EGFR DN (gray). N=14 for control branches; P55, N=16 for EGFR DN branches; P55, N=14 for control branches; P65, N=11 for EGFR DN branches; P65, N=16 for control branches; P72, N=18 for EGFR DN branches, N=18 branches for control branches; adult, N=22 branches for EGFR DN branches; adult. Kruskal–Wallis and Dunn’s as post-hoc test; \*\*\*\**p* < 0.0001. **(O-P)** Labelling of post-synaptic partners (magenta) of adult DCNs (green) using Trans-Tango at 25°C for 7 days showing reduced connectivity in EGFR DN **(P)** compared to control **(O). (Q)** Quantification showing reduced post-synaptic cell body number in medulla in control (black) vs EGFR DN (gray) in adult DCNs at 25°C for 7 days old adults. N=14 optic lobes for control and N=14 optic lobes for EGFR DN; \*\*\*\**p* < 0.0001, Mann Whitney test. **(R/R’-T/T’)** Rescue of BrpD3 GFP (green) puncta and secondary branches in adult DCNs labeled by uas-CD4 tomato (magenta) with increased gene copy number of Brp (Pacman Brp) in EGFR DN background **(T/T’)** back to control level **(R/R’)** compared to EGFR DN alone **(S/S’)** in adults. **(U)** Adult quantification showing the rescue of BrpD3 GFP puncta normalized to individual axon branch length in Pacman Brp + EGFR^DN^ (orange) back to control (black) level compared to EGFR DN (gray). N=58 primary branches for control, N=30 primary branches for EGFR DN and N=56 primary branches for Pacman Brp+ EGFR DN. Kruskal– Wallis and Dunn’s as post-hoc test; \*\*\*\**p* < 0.0001. **(V)** Quantification showing no rescue of adult primary branch number per axon in Pacman Brp + EGFR DN (orange) back compared to EGFR DN (gray). N=18 axons for control, N=24 axons for EGFR DN and N=26 axons for Pacman Brp+ EGFR DN. Kruskal–Wallis and Dunn’s as post-hoc test; \*\*\*\**p* < 0.0001. **(W)** Quantification showing rescue of adult secondary branch number per primary branch in Pacman Brp + EGFR DN (orange) back to control (black) level compared to EGFR DN (gray). N=20 primary branches for control, EGFR DN and Pacman Brp + EGFR DN. Kruskal–Wallis and Dunn’s as post-hoc test; \*\**p* = 0.0007. Error bars denote mean±SEM, scalebar represents 5μm except **(O**,**P)** where it represents 20μm.

Given that EGFR and Brp enter branches with different temporal dynamics (EGFR before Brp), we asked when EGFR is required for the presence of Brp in branches by examining the recruitment and accumulation of BrpD3 GFP in M-DCN axons and branches over time. Inactivation of EGFR did not interfere with the initial recruitment of BrpD3 at P55 or at P65 where the increase in the number of branches caused by EGFR inactivation also caused an increase in the number of BrpD3 puncta. In contrast, we observed a significant decrease of BrpD3 levels starting at P72 and continuing into adults (Fig.3H,H’-M,M’,N). To examine the temporal dynamics of the gradual loss of BrpD after initial recruitment, we performed live imaging and calculated the stability of BrpD3 GFP puncta in branches during synaptogenesis. We tracked the stability of Brp puncta for 2 hours in our *ex-vivo* cultures at P65 and P72 and classified puncta as stable if they were present throughout tracking, or as unstable if they disappeared at any point during the tracking. Between P65 and P67, in control DCNs, 82% of Brp puncta were stable, compared to 57% in EGFR-DN DCNs. Between P72 and P74, 93% of Brp puncta became stable in controls compared to 76% in EGFR DN expressing DCNs (Fig.S3G, Supplemental Video S3). Because Brp acts as a scaffold protein for active zone formation (Huang *et al*., 2020), these observations suggest that lack of EGFR activity reduces the stability of active zones during late brain development.

Altogether, the data above show that while EGFR inactivation causes an increase in primary and secondary axon branches, it causes a sharp decrease in the presynaptic active zone protein Brp. What is the impact of these seemingly opposing changes on DCN postsynaptic connectivity and circuit wiring? To test this, we first needed to determine the connectome of DCNs in the medulla. We used the anterograde trans-synaptic method for target tracing approach “Trans-Tango”, which labels all the post-synaptic targets in an unbiased manner without required prior knowledge of cell types (Talay *et al*., 2017). We used stringent conditions (see methods) to optimize sparse labelling of postsynaptic targets (Fig.S4A-D). We found that M-DCNs connect to a large variety of medulla projecting neurons along with some lobula and lobula plate targeting neurons (Fig.S4F1-F21). Their most frequent partners are lamina wide field cells (Lawf1/2) (Fig.S4F18,G), followed distantly by trans medulla neurons (Tm2/21/Y8/9) (Fig.S4F2,F3,F7,F8,G) (Fischbach and Dittrich, 1989). We further validated Lawf1 and Tm2 subtypes using activity dependent GRASP which confirmed them as DCN postsynaptic targets (Fig.S4H-H’’,I-I”). Next, we tested the effects of loss of EGFR activity on DCN circuit wiring. We observed a drastic reduction in overall connectivity (Fig.3O-Q, Fig.S4C-E), with a significant reduction in the most frequent partners, Lawf1/2, Tm2/21, and a complete loss of less frequent partners (Fig.S4G,G’). Therefore, the stabilization of Brp by EGFR is required for the specific pattern of M-DCN connectivity.

Thus far we have found that knock-down of Brp causes an increase in secondary branching and that loss of EGFR activity results in a late loss of Brp puncta as well as a similar increase in secondary branching (Fig.S2G, Supplemental Video 2.1). We therefore asked if reduction of Brp might explain the secondary branching phenotype of EGFR inactivation. To test this idea, we introduced an extra genomic copy of the *brp* gene in a DCN EGFR-DN background. Remarkably, increasing Brp levels with one genomic copy rescues the loss of BrpD3-GFP puncta showing specificity of the phenotype (Fig.3R-T,U). The extra copy of *brp* also rescued the secondary branching phenotype (Fig.3R-T,W) but did not rescue the primary branching phenotype associated with the early role of EGFR (Fig.3R-T,V). Together these data show that inactivation of EGFR causes loss of Brp which in turn destabilizes terminal axon branches leading to an increase in their numbers. We conclude that the quantity of stable Brp is a determinant of terminal branch stability.

### Temporally specific requirement for EGFR activity in synapse formation and terminal branching dynamics

We have previously shown that early DCN branch pruning is regulated by EFGR via the control of actin dynamics. In this study, we have observed that EGFR is also required during a second phase to maintain Brp and synaptic connectivity. We asked whether these two effects reflect different temporal requirements of EGFR, or whether the late phenotype is an indirect consequence of the early effect of EGFR on primary branching. To distinguish between these two possibilities, we used temperature sensitive inactivation of EGFR, by inducing the EGFR-DN transgene at different times during development using the Gal4 temperature-sensitive repressor Gal80^ts^. We ascertained that the transgene is active and prevents GFP expression at 22°C (Fig.S5A) and calibrated the temporal progression of branch development at 22°C with respect to control at 25°C (Fig.S5B-F). Gal80^ts^ prevents EGFR-DN expression (and thus allows EGFR activity) at 22°C (green bar). At 29 degrees Gal80^ts^ is inactivated, thus allowing EGFR-DN to be expressed, which in turn will inhibit EGFR activity (magenta bar). Thus, at 22°C EGFR is active, while at 29°C EGFR is inactive. As expected, continuously inactivating EGFR starting at P48 till adult results in a significant increase in primary and secondary branches (asterisks) and a significant decrease in Brp (Fig.4A-E). Inactivating EGFR only early between P42 and P57 increases primary branches but has no effect on secondary branches (asterisks) or Brp (Fig.4F-J). In contrast, inactivating EGFR starting either at P55 (Fig.4K-O) or at P65 (Fig.4P-T) has no effect on primary branch number, but significantly increases secondary branches (asterisks) and significantly decreases Brp. This role of EGFR in synapse formation appears strictly developmental as inactivating EGFR specifically in adults does not result in any defects in branching or Brp levels and distribution (Fig.S5G-K). Therefore, mechanistic regulation of EGFR at pre-synaptic branches defines a developmental critical interval for terminal branch consolidation and synapse stabilization required for neuronal circuit wiring in adults.

**Fig.4.**
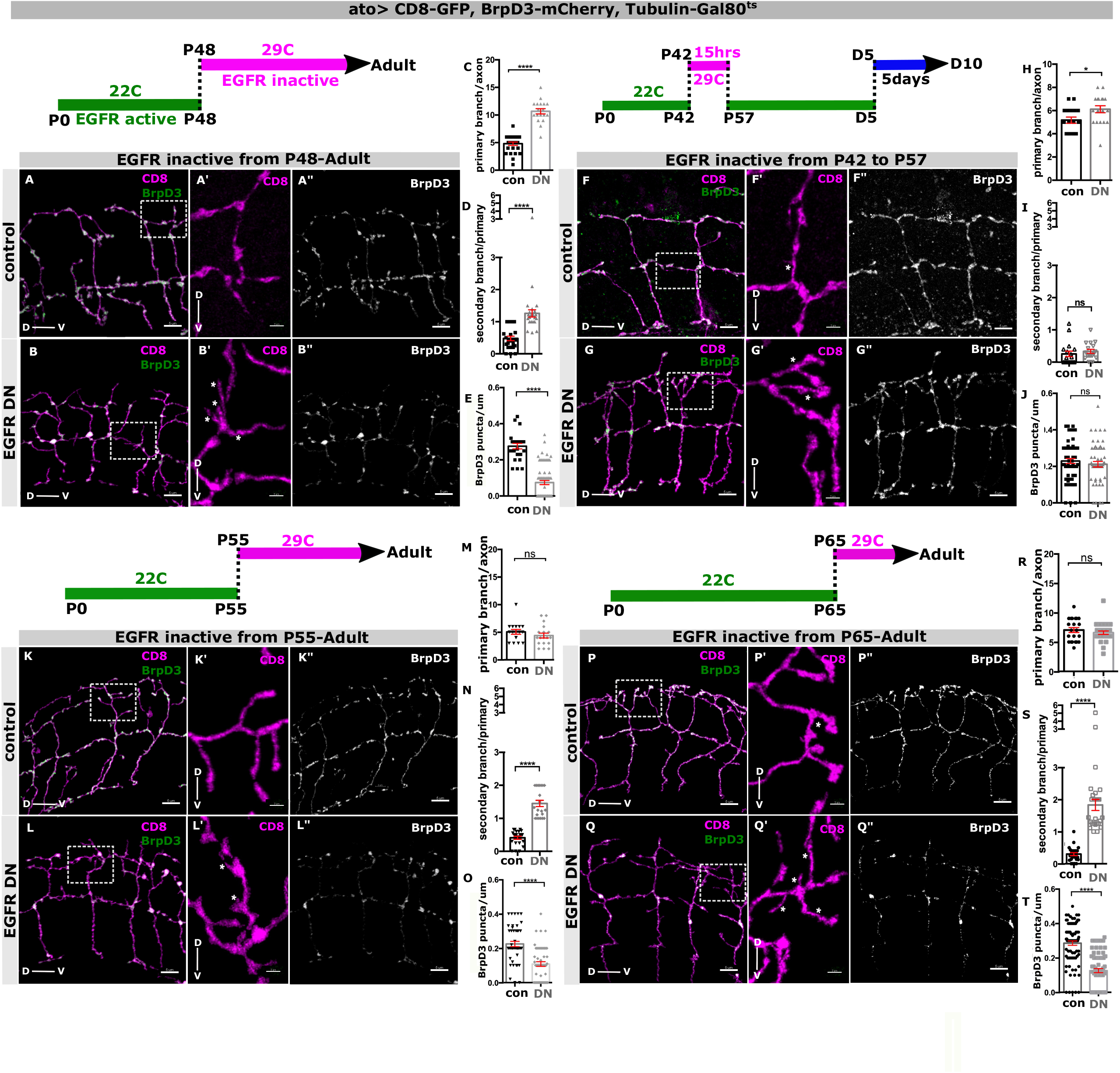
Temporally specific requirement for EGFR activity in synapse formation and primary branch consolidation: Adult DCN branch morphology labeled by uas-CD8 GFP (green) and AZs marked by uas-BrpD3-mCherry (red). EGFR is active when shifted to 22°C (green bar) whereas inactive when shifted to 29°C (magenta bar). We inactivated EGFR temporally by shifting the Gal80^ts^ construct to 29°C during different developmental time intervals and analyzed young adults unless otherwise stated. White squares represent regions of higher magnification. **(A/A”-B/B”)** Lack of EGFR activity from P48 throughout development results in more primary and secondary branching (asterisks) with reduced BrpD3-mCherry puncta in adult DCNs compared to control. **(C)** Quantification showing increased primary branch number per axon in control (black) vs EGFR DN (gray). N=25 axons for control, N=26 axons for EGFR DN. Mann Whitney test; \*\*\*\**p* = 0.0001. **(D)** Quantification showing increased secondary branches per primary branch in control (black) vs EGFR DN (gray). N=25 primary branches for control and N=22 primary branches for EGFR DN. Mann-Whitney test; \*\*\*\**p* < 0.0001. **(E)** Quantification showing decreased BrpD3-mCherry puncta per unit branch length in control (black) vs EGFR DN (gray). N=25 branches for control and N=66 branches for EGFR DN. Mann Whitney test; \*\*\*\**p* = 0.0001. **(F/F”-G/G”)** Lack of EGFR activity only between P42 to P57 during development results in more primary branching with no effect on secondary branching (asterisks) or BrpD3-mCherry puncta localisation in adult DCNs compared to control. The young adults were shifted to 25°C for 5 days after eclosion for transgene expression prior to analyses. **(H)** Quantification showing increased primary branch number per axon in control (black) vs EGFR DN (gray). N=17 axons for control and N=18 axons for EGFR DN. Mann-Whitney test; \**p* =0.0139. **(I)** Quantification showing unaffected secondary branch number per primary branch in control (black) vs EGFR DN (gray). N=17 primary branches for control and EGFR DN. Mann-Whitney test; ^**ns**^p=0.1462. **(J)** Quantification showing comparable BrpD3 mCherry puncta per unit branch length in control (black) vs EGFR DN (gray). N=59 primary branches for control and N=56 primary branches for EGFR DN. Mann-Whitney test; ^**ns**^p=0.36. EGFR activity blocked only late; either from P55 **(K/K”-L/L”)** or from P65 **(P/P”-Q/Q”)** throughout development leads to decreased BrpD3-mCherry puncta and increased secondary branches (asterisks) with no significant change in primary branches in adult DCNs. **(M**,**R)** Quantification showing unaffected primary branches per axon in control (black) vs EGFR DN (gray). **(M)** N=38 axons for control, N=43 axons for EGFR DN. Mann Whitney test; ^**ns**^*p*=0538. **(R)** N=36 axons for control, N=26 axons for EGFR DN. Mann Whitney test; ^**ns**^*p*=7204 **(N**,**S)** Quantification showing increased secondary branches per primary branch in control (black) vs EGFR DN (gray). **(N)** N=22 primary branches for control and N=21 primary branches for EGFR DN. Mann-Whitney test; \*\*\*\**p* < 0.0001. **(S)** N=25 primary branches for control and n=26 primary branches for EGFR DN. Mann-Whitney test; \*\*\*\**p* < 0.0001. **(O**,**T)** Quantification showing decreased BrpD3-mCherry puncta per unit branch length in control (black) vs EGFR DN (gray). **(O)** N=47 primary branches for control and N=53 primary branches for EGFR DN. Mann Whitney test; \*\*\*\**p* < 0.0001. **(T)** N=87 primary branches for control and N=94 primary branches for EGFR DN. Mann Whitney test; \*\*\*\**p* < 0.0001. Error bars denote mean±SEM, scalebar represents 5μm.

### EGFR activity is required to prevent Brp degradation during synaptogenesis: (fig.5, S6)

We have previously shown that autophagy can regulate synapse formation by restricting filopodial kinetics (Kiral *et al*., 2020) and EGFR has been shown to regulate autophagy in *Drosophila* testis (Sênos Demarco and Jones, 2020) and in many tumorigenic contexts (Wu and Zhang, 2020). We asked whether loss of EGFR activity results in increased Brp degradation. To this end, we first used a general degradation reporter (myr-mCherry-pHluorin) (Jin *et al*., 2018) and observed a significant increase in localization of this probe in acidic compartments (Fig.S6A-C). Next, we examined the proportion of BrpD3-GFP puncta in acidic late endosomal/autophagosomal compartments in control and EGFR-DN M-DCNs using endogenous Rab7, DCN expressed Rab7-RFP and DCN-expressed pH sensitive BrpD3-mCherry-pHluorin, as independent markers of such compartments. We observed a progressive increase in the colocalization of BrpD3-GFP with endogenous Rab7 in axon branches upon EGFR inactivation starting at P65 and continuing into adults (Fig.5A-F,G). We obtained similar results using the Rab7-RFP reporter expressed specifically in DCNs (Fig.S6D-G). Importantly, this colocalization was not observed for Syd1 (Fig.S6H-J), whose levels and distribution were not affected by EGFR inactivation (Fig.3B,B’,E). Finally, the BrpD3-mCherry-pHluorin probe showed increased localization into acidic compartments upon EGFR inactivation as detected by increased mcherry to pHluorin signal intensity starting at P72 and into adults (Fig.5H-M,N).

**Fig.5.**
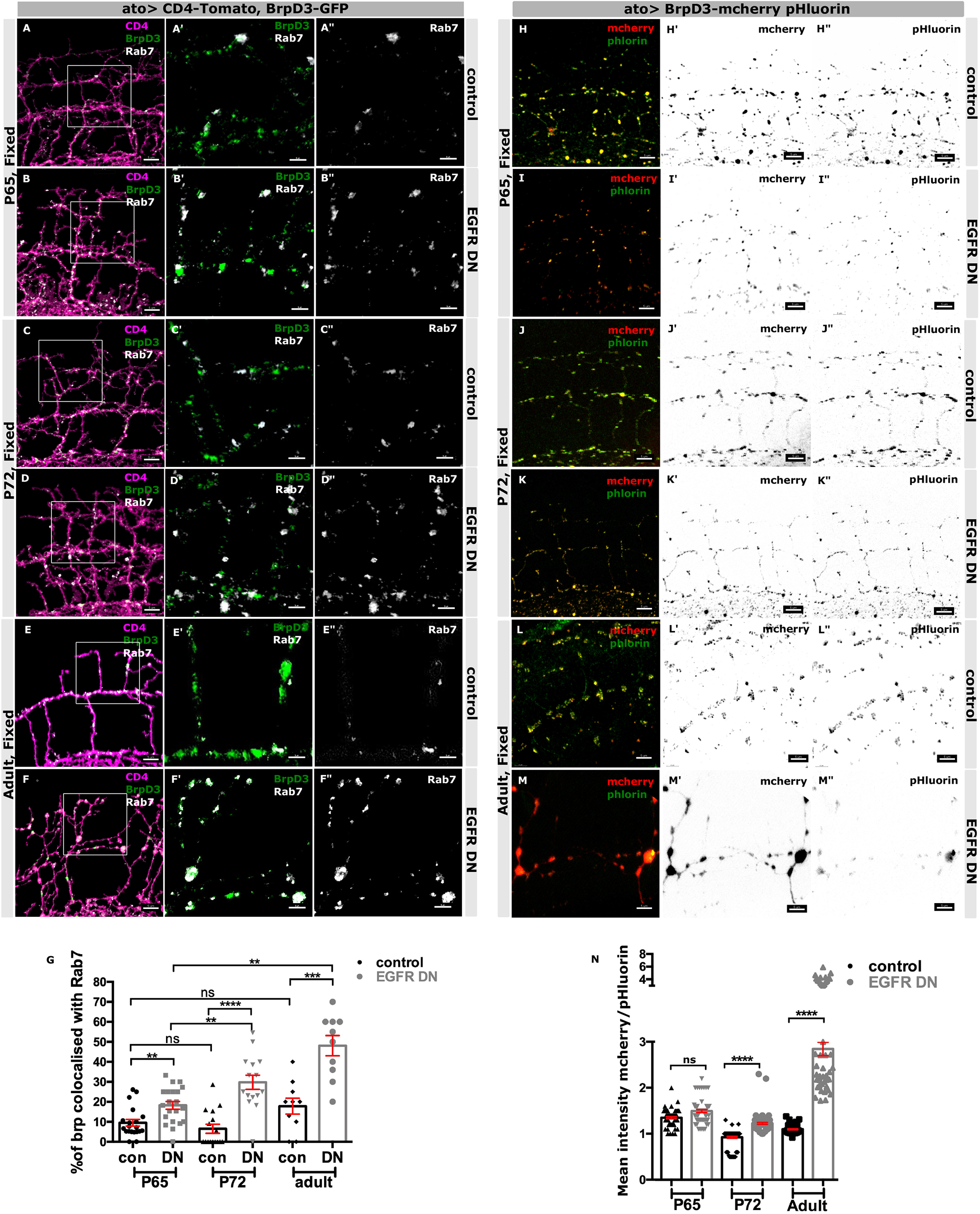
EGFR activity is required to prevent Brp degradation during synaptogenesis: **(A/A”-F/F”)** Progressively increased colocalization of endogenous Rab7 (white) with BrpD3-GFP puncta (green) in DCN axon terminals labeled with CD4-tomato (magenta) in control vs EGFR DN during late development; P65 **(A/A”-B/B”)**, P72 **(C/C”-D/D”)** and adult **(E/E”-F/F”)**. White squares represent regions of higher magnification showing Rab7-BrpD3 colocalization in **(A’/A”-F/F”). (G)** Quantification showing increased percentage of BrpD3-GFP puncta colocalized with endogenous Rab7 per axon at P65, P72 and adults in control (black) vs EGFR^DN^ (gray). N=20 axons for control vs N=21 axons for EGFR DN at P65;, N=16 axons for control vs N=15 axons for EGFR DN at P72, N=10 axons for control vs N=10 axons for EGFR DN at adults. \*\*\*\**p* < 0.0001, Kruskal Wallis and Dunn’s as post-hoc test. **(H/H”-M/M”)** Simultaneous increase in mCherry signal (red) intensity compared to pHluorin (green) intensity when expressing the degradation probe uas-BrpD3-mCherry-pHluorin in DCN branches in EGFR DN compared to control during late development: P65 **(H’/H”-I/I”)**, P72 **(J/J”-K/K”)** and adult **(L/L”-M/M”). (N)** Quantification showing increased mean intensity of mCherrry channel to pHluorin channel across several BrpD3-DF puncta per axon during P65, P72 and adults in control (black) vs EGFR DN (gray). N=65 BrpD3-DF punctum for control vs N=65 BrpD3-DF punctum for EGFR DN at P65, N=82 BrpD3-DF punctum for control vs N=83 BrpD3-DF punctum for EGFR DN at P72; N=68 BrpD3-DF puncta for control vs N=47 BrpD3-DF puncta for EGFR DN at adults. \*\*\*\**p* < 0.0001; Kruskal Wallis and Dunn’s as post-hoc test; Error bars denote mean±SEM, scalebar represents 5μm; apart from **(A’/A”, B’/B”, C’/C”, D’/D”, E’/E”, F’**,**F”)** which represents 3μm.

These data suggest that EGFR is not required for increasing the levels of Brp import into the synaptic terminal, but instead for maintaining Brp pools already present in presynaptic branches by preventing their degradation. If Brp degradation is already fully suppressed by wild type levels of EGFR activity, then increasing EGFR activity further should not lead to an increase in Brp levels. We tested this idea by examining Brp upon expression of a constitutively activated EGFR (EGFR-CA) in DCNs. We found no effect on BrpD3 density or distribution, nor on the degree of colocalization with Rab7-RFP (Fig.S6K-N).

### EGFR signals through autophagy to maintain DCN presynaptic active zone, circuit connectivity, and behavior: (fig.6, fig.7)

To investigate the potential causal role of increased degradation for the loss of Brp and synaptic connectivity, we performed knockdown of the autophagic regulators Rab7 and Atg6 upon inhibition of EGFR activity and assayed terminal branching, BrpD3 distribution, and M-DCN postsynaptic connectivity. Knockdown of either Rab7 or Atg6 alone did not cause a significant change in the number of terminal secondary branches or level of BrpD3 (Fig.6A,A’-C,C’,M,N). In contrast, knockdown of either Rab7 or Atg6 upon EGFR inactivation completely suppressed the increase in secondary branches and restored Brp at M-DCN synaptic terminals back to control levels (Fig.6D,D’-F,F’,M,N), demonstrating that autophagy is required for the synaptic loss caused by EGFR inactivation.

**Fig.6.**
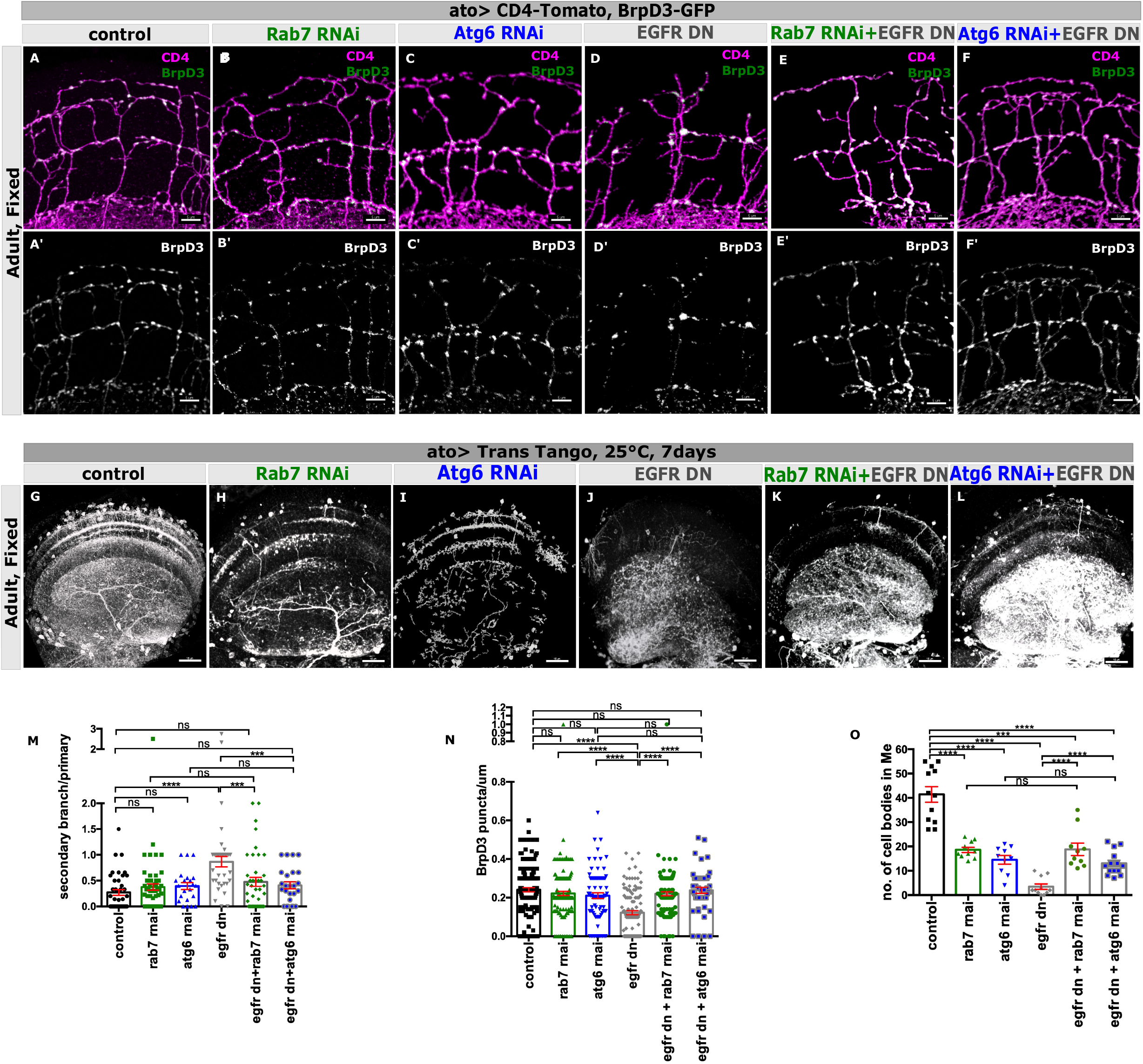
EGFR signals through autophagy to maintain DCN presynaptic active zone and circuit connectivity: **(A/A’-F/F’)** Rescuing loss of adult DCN terminal axon branch (magenta) number and BrpD3-GFP puncta (green) distribution in Rab7 RNAi + EGFR DN **(E/E’)** and Atg6 RNAi + EGFR DN **(F/F’)** back to the level of control **(A/A’)**, Rab7-RNAi **(B/B’)** or Atg6-RNAi **(C/C’)** as compared to EGFR DN **(D/D’). (G-L)** Corresponding rescue of post-synaptic partner (white) loss in DCNs using Trans-Tango at 25°C for 7 days old adults in EGFR DN **(H)** by expressing Rab7 RNAi + EGFR DN **(K)** or Atg6 RNAi + EGFR DN **(L)** back to the level of Rab7-RNAi **(H)** or Atg6-RNAi **(I)** respectively but only partial rescue compared to control **(G). (M)** Quantification showing rescue of secondary branch number per primary in Rab7 RNAi + EGFR DN (green+gray) and Atg6 RNAi + EGFR DN (blue+gray) back to the level of control (black), Rab7 RNAi (green), Atg6 RNAi (blue) as compared to EGFR DN (gray). N=41 branches for control, N=50 branches for Rab7 RNAi, N=22 branches for Atg6 RNAi, N=35 branches for EGFR DN, N=41 branches for Rab7 RNAi + EGFR DN, N= 25 branches for Atg6 RNAi + EGFR DN. Kruskal–Wallis and Dunn’s as post-hoc test; \*\*\*\**p* < 0.0001. **(N)** Quantification showing similar rescue of BrpD3-GFP puncta per unit branch length in Rab7 RNAi + EGFR DN and Atg6 RNAi + EGFR DN back to the level of control, Rab7 RNAi, Atg6 RNAi as compared to EGFR DN. N=153 branches for control, N=138 branches for Rab7 RNAi, N=84 branches for Atg6 RNAi, N=124 branches for EGFR DN, N=122 branches for Rab7 RNAi + EGFR DN, N=50 for Atg6 RNAi + EGFR DN; Kruskal–Wallis and Dunn’s as post-hoc test; \*\*\*\**p* < 0.0001. **(O)** Quantification for rescuing post-synaptic cell body number in medulla per optic lobe in Rab7 RNAi + EGFR DN and Atg6 RNAi + EGFR DN back to the level of control, Rab7 RNAi, Atg6 RNAi as compared to EGFR DN. N=12 optic lobes in control, N=11 optic lobes in Rab7 RNAi, N=10 optic lobes in Atg6 RNAi, N=13 optic lobes in EGFR DN, N=12 optic lobes in Rab7 RNAi + EGFR DN and N=14 optic lobes in Atg6 RNAi + EGFR DN. Kruskal–Wallis and Dunn’s as post-hoc test; \*\*\*\**p* < 0.0001. Error bars denote mean±SEM, scalebar represents 5μm except **(G-L)** where it is 15μm.

Next, we asked whether this cell-autonomous rescue of presynaptic terminals suffices to restore M-DCN postsynaptic connectivity. Downregulation of either Rab7 or Atg6 alone resulted in a ∼50% decrease in the number of trans-tango labelled M-DCN postsynaptic cells, while EGFR inactivation caused an almost complete loss of connectivity (Fig.6G-J,O). Importantly however, knockdown of either Rab7 or Atg6 in the EGFR-DN background rescued the connectivity to postsynaptic cells back to the levels observed upon knockdown of Rab7 or Atg6 alone (Fig.6J-L,O) showing that the effect of EGFR inactivation requires autophagy.

We have previously shown that the M-DCNs regulates behavioral responses in the multiparametric single fly visual response assay called Buridan’s paradigm (Linneweber *et al*., 2020). In this assay, single flies walk freely between two identical visual cues (Fig.7A) and reduction of synapses causes increased fly activity when flies were tested at the same temperature at which they developed (Kiral *et al*., 2021). Consistent with this, we find that inactivation of EGFR specifically in DCNs results in flies walking longer distances, staying active for a longer time, and increasing the number of walks between the two cues (Fig.7B-H; Table S1). Specifically, we observed ∼1.12-fold increased distance travelled (total path length between the 2-stripes (Fig.7F), ∼1.14 fold increased activity time (total time the flies were moving) (Fig.7G) and ∼1.2 fold more number of walks (Fig.7H). Remarkably, DCN-specific knockdown of Rab7 in in the EGFR-DN background completely rescued all these phenotypes back to control levels (Fig.7D,F-H), suggesting that even partial rescue of DCN synaptic connectivity (Fig.6O), is sufficient to support normal behavioral activity at the level assayed here.

**Fig.7.**
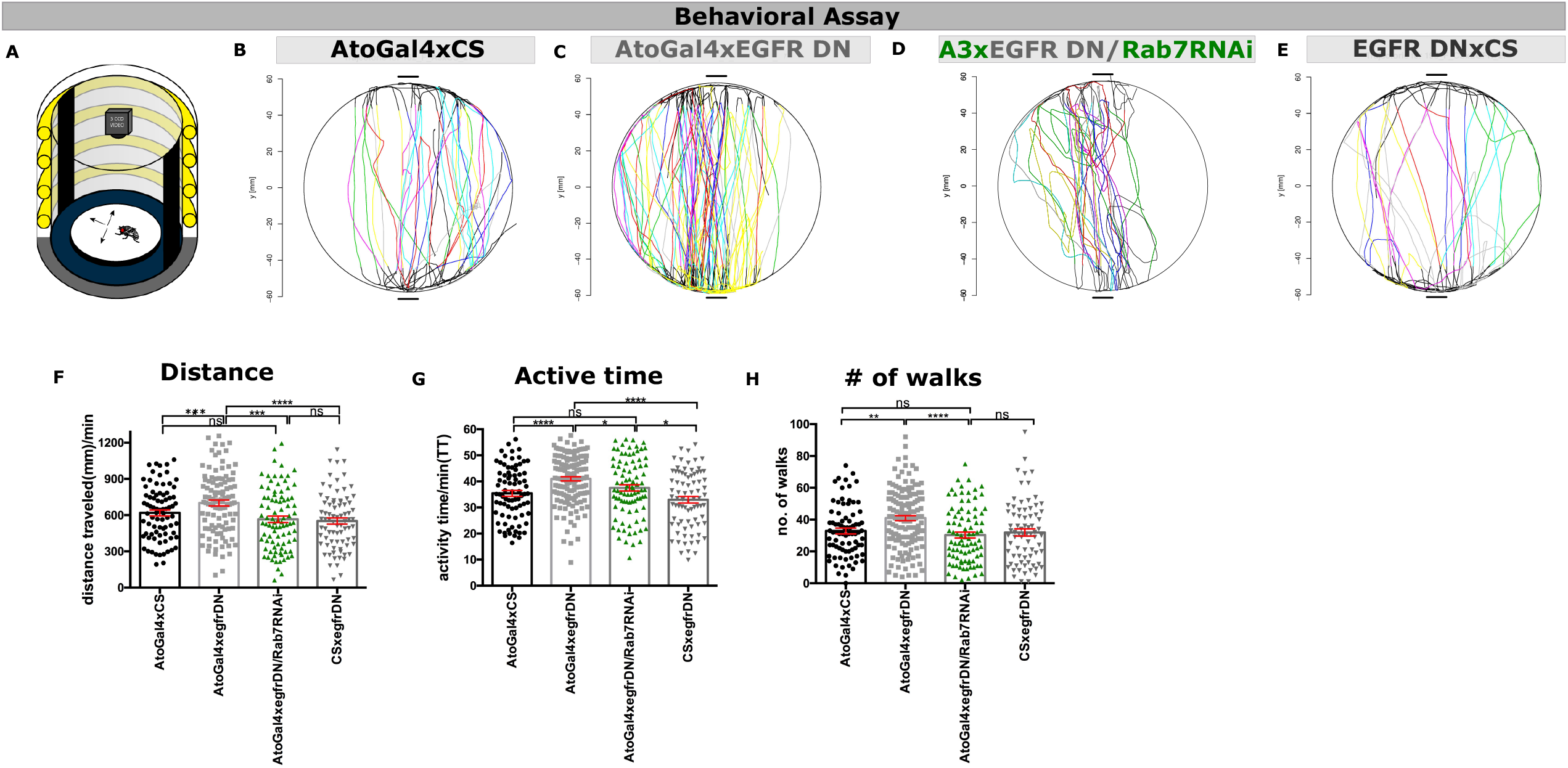
**(A)** Schematic illustration of the adult fly behavior assay; Buridan’s paradigm. **(B-E)** Individual fly walking tracks between 2 black stripes in AtoGal4-14a(A3)xCS**(B)**, Atogal4-14a(A3)/EGFR DN **(C)** Atogal4-14a(A3)/ EGFR DN+ Rab7 RNAi **(D)** EGFR DNxCS **(E). (F-H)** Quantification of fly activity behavior using 3 parameters: distance traveled **(F)**, activity time **(G)** and number of walks **(H)** showing rescue in behavioral phenotype in Atogal4-14a/ EGFR DN+ Rab7 RNAi (gray+green) back to the level of AtoGal4-14axCS (black) or EGFR DNxCS (dark gray) compared to Atogal4-14a(A3)/EGFR DN (gray). Each dot represents individual flies. N=80 individuals for AtoGal4xCS, N=106 individuals for AtoGal4x EGFR DN, N=89 individuals for AtoGal4x EGFR DN/Rab7 RNAi, N=79 individuals for CSxEGFR DN. Kruskal–Wallis and Dunn’s as post-hoc test; \*\*\**p* = 0.0002 **(F)**, \*\*\**p* = 0.0005 **(G)**, \*\*\*\**p* < 0.0001 **(H)**. Error bars denote mean±SEM.

## Discussion

The complexity of neuronal circuit wiring patterns, driven to a significant extent by the degree of axonal branching during development (Hoersting and Schmucker, 2021), is thought to be key to the emergence of complex behaviors and cognitive capacities. In the mammalian motor cortex for example, axon branch complexity can allow a single neuron to innervate very distant ipsi- and contra-lateral cortical and sub-cortical targets (Economo *et al*., 2016). Some genetic risk factors for human neuropsychiatric disorders such as Schizophrenia appear to be associated with alterations in terminal axonal branches (Shao *et al*., 2019). In insects, the relative conservation of axonal projections patterns is thought to underlie species-specific innate behavioral patterns while permitting individual variation in these behaviors. For instance, the unique branching pattern of L-fibers supports the nocturnal lifestyle of *M*.*genalis* (Grueber *et al*., 2005), while intrinsic variation in M-DCN connectivity underlies individual variation in *Drosophila* visual response behavior (Linneweber *et al*., 2020). While it is critical to dissect the specific phenomena of axonal branching and synaptogenesis during circuit wiring, it is equally critical to understand the spatial, temporal, and mechanistic coordination of these processes during development to arrive at a more complete description of the emergence of both the conserved patterns and the individual variation of neuronal circuit diagrams.

Here we asked if, when and how molecular factors that play key roles in regulating axon branching, synapse formation and membrane degradation interact to produce specific presynaptic patterns, neuronal circuit connectivity and behavior using the developing *Drosophila* visual system as a model. This diverse list of cellular processes is tied together by their collaborative effects during the development of synaptic connectivity. Axo-dendritic branching and synapses formation have long been known to depend on each other during synaptotropic growth of branches based on synapse stabilization (Vaughn, Barber and Sims, 1988). The developmental interactions of axonal and dendritic processes are a major contributor, and can sometimes predict, the adult synaptic connectivity (Agi, Kulkarni and Hiesinger, 2020). By contrast, cell biological processes like cytoskeletal dynamics or membrane degradation have long been described as a ‘permissive’ basis for more ‘instructive’ molecular mechanisms of synaptic specification. However, to the extent that such cell biological mechanisms directly contribute to branching and synapse formation, they become parts of a composite instruction that cannot be pinned on a single molecular mechanism but require the consideration of several collaborating factors to understand a neuron’s choice to branch and form a synapse (Hiesinger, 2021). Hence, an integrative analysis of molecular recognition, signaling and cell biological machinery is necessary to mechanistically understand how branching contributes to adult synaptic connectivity.

Numerous molecular and cellular mechanism have been implicated in the spatiotemporal control of how synaptic partners are brought together for synapse formation (Agi, Kulkarni and Hiesinger, 2020). Over the last 25 years, investigating the role of EGFR in nervous systems has demonstrated its roles in neural stem cell maintenance (Aguirre, Rubio and Gallo, 2010), astrocyte and oligodendrocyte maturation (Galvez-Contreras, Quiñones-Hinojosa and Gonzalez-Perez, 2013), axon regeneration (Koprivica *et al*., 2005) and more recently in neurite outgrowth and branching (Goldshmit *et al*., 2004). In addition, we have recently demonstrated roles for the cell biological regulation of filopodial dynamics and autophagic degradation in establishing specific synaptic connectivity (M Neset Özel *et al*., 2019; Kiral *et al*., 2020). How such basic cellular processes are coupled to the process of patterning axonal branches and how the three processes – branching, synaptogenesis and autophagy – are coordinated in time to establish circuit-specific wiring diagrams has remained unclear. We find that that these processes are indeed coupled but only during a very specific temporal interval through the activity of the Epidermal Growth Factor Receptor (EGFR) preventing the autophagic degradation of the synaptic active zone protein Bruchpilot (Brp). Inactivation of EGFR during this specific critical interval in late development, but not before or after, causes Brp degradation, changes to circuit wiring and altered visually driven behaviors.

An interesting observation are the differences in dynamics upon inactivation of EGFR and knockdown of Brp. While both result in increased secondary branches, loss of Brp leads to a more dynamic state of these branches even in the adult brain. In contrast, loss of EGFR causes these secondary branches to be largely static, even though EGFR inactivation also causes Brp loss. This is likely because EGFR loss also impairs actin dynamics which are required for filopodial dynamics, while Brp loss does not affect EGFR activity and thus does not impair actin dynamics. These observations underline the critical importance of live imaging for discovering and interpreting otherwise seemingly similar phenotypes. These observations are also consistent with the genetic hierarchy we observed whereby EGFR regulates BrpD3 levels, but Brp does not appear to regulate EGFR levels. Again, live imaging combined with temporally restricted inactivation of EGFR allowed us to dissect the temporal logic of this genetic hierarchy and discover the role of EGFR activity as coupling mechanism between branching dynamics, local degradation, and synapse formation.

Our observations show that the coupling between branching, autophagy and synaptogenesis is only required during a very specific temporal interval which, perhaps not surprisingly, coincides with active synaptogenesis in the developing fly brain [REF]. Our analysis of the temporal sequence of trafficking of various molecules indicates that this developmental critical interval is opened by the spatial and temporal coincidence of EGFR and BrpD3 in exploratory branches. Similarly, it is likely that the critical interval ends once synaptic contacts between DCNs and their postsynaptic cells have been established. What remains to be explored is how exactly synapse formation ends the interactive feedback between branch dynamics and synapse degradation. One possibility is that postsynaptic dendrites provide molecular signals to presynaptic axonal branches to limit active zone degradation. Another possibility is that the initiation of spontaneous activity alters local endolysosomal recycling (Tagliatti *et al*., 2016) to reduce degradation and favor maintenance, for example through the activity-dependent regulation of local translation of resident mRNAs (Rajgor, Welle and Smith, 2021). A third possibility is the late arrival of presynaptic proteins that protect the active zone, reduce autophagy or alter EGFR function. Clearly, none of these mechanisms are mutually exclusive and we speculate that a combination of such feedback mechanisms would be the best way to ensure a robust closing of the developmental critical interval we discovered here. Flies, like vertebrates, undergo a critical period of postnatal experience-dependent synaptic pruning with unknown molecular mechanism. It would be interesting to explore whether the mechanism we uncovered here also plays a role in synaptic regulation and neural circuit plasticity during that critical period. It is possible that similar molecular modules are used re-iteratively to regulate synaptic homeostasis and pre-synaptic branching in experience-independent and dependent brain development.

We have previously shown that an increase in local autophagy also led to a decrease of adult synapses in R7 photoreceptor neurons (Kiral *et al*., 2020), albeit by a different mechanism. In R7 photoreceptors, local autophagosome formation at the tips of synaptogenic filopodia is accompanied by engulfment of synaptic seeding factors but not Brp, followed by filopodial collapse and thus less filopodial availability to form synapses. In DCNs, axonally localized autophagosome formation is accompanied by engulfment of Brp but not synaptic seeding factors, leading to a destabilization and reduction of mature synapses. Hence, the spatiotemporal specific roles of autophagy in different types of axon terminals during synapse formation are highly context-specific, involve different substrates, and yet lead to similar outcomes. An obvious contextual difference between R7s and DCNs is that only the latter form branched axons and employ synaptotropic-like branch stabilization through mature synapses.

Increasing evidence supports the important roles that stochastic and/or noisy molecular processes play in the early phases of establishing wiring specificity prior to the final step of spatially and temporally restricted local synaptic matching (Hassan and Hiesinger, 2015). While neuronal circuit wiring patterns across the animal kingdom are highly robust, they, almost without exception, show at least some degree of intrinsic variation within and between individuals. Noise in the genetically encoded program can ensure robustness of highly reproducible wiring patterns, for example when stochastic exploration ensures partner finding (Hiesinger and Hassan, 2018). Correspondingly, axon branch initiation in DCNs is noisy and exploratory, while the final axonal pattern is robustly stereotypic. The evidence presented in this work demonstrates the importance of intrinsic local control of the feedback between axonal branching and synapse formation to ensure the robust outcome. Such local feedback regulation ensures that genetically encoded noisy molecular and cellular processes such as filopodial growth and retraction and synaptic seeding are coordinated in time and space to produce conserved, robust, yet individually variable, non-random neuronal circuit diagrams.

## ACKNOWLEDGMENTS

This work was supported by the Einstein-BIH program and DFG Research Unit 5289 RobustCircuit project P2 (to B.A.H and P.R.H), DFG Research Unit Syntophagy RP7 (Hi 1886/8), funding from the European Research Council (ERC) under the European Union’s Horizon 2020 research and innovation programme (grant agreement No. 101019191 (to P.R.H), the Investissements d’Avenir program (ANR-10-IAIHU-06), Paris Brain Institute-ICM core funding, the Paul G. Allen Frontiers Group Allen Distinguished Investigator grant, the Roger De Spoelberch Prize and an NIH Brain Initiative RO1 grant (1R01NS121874-01) (to B.A.H.). M.A. is funded by the Fondation de la Recherche Medical (FRM) postdoctoral fellowship (ARF202005011913). We thank the BioSupraMol Optical Microscopy facility for STED microscopy, members of the Hassan and Hiesinger labs for helpful discussions and Drs. Stephan Sigrist and Dietmar Schmucker for comments on the manuscript.

## AUTHOR CONTRIBUTIONS

S.D., B.A.H. and P.R.H. conceived the study, designed the experiments, and wrote the manuscript. S.D., G.A.L. and M.A. conducted all experiments and data analysis.

## Materials and Methods

### Experimental model and subject details

Flies were reared at 25°C on standard cornmeal/yeast diet for all crosses and at 21°C and 29°C for Gal80ts experiments. For developmental analyses, white pre-pupae (P+0%) were collected and incubated at 25°C to pupal stages as stated on figures. The following *Drosophila* strains were either obtained from Bloomington *Drosophila* Stock Center (BDSC), Vienna *Drosophila* Resource Center (VDRC) or other groups: ato-Gal4-14a (Hassan Bassem A. et al., 2000), M DCN-Gal4 (Linneweber *et al*., 2020), ato-LexA (Langen *et al*., 2013), Lawf1-Gal4 (Konstantinides *et al*., 2018); Tm2-Gal4 (Ting *et al*., 2011), nsyb-GRASP flies (BDSC), UAS-Denmark (BDSC), UAS-CD4 tdGFP(BDSC), UAS-CD4 tdTomato(BDSC), UAS-CD8 GFP(BDSC), UAS-BrpD3 GFP (Schmid *et al*., 2008), UAS-BrpD3 mCherry (Schmid *et al*., 2008), UAS-Syd1 GFP (Owald *et al*., 2010), UAS-EGFR DN (Zschätzsch *et al*., 2014), UAS-Rab7 RFP(BDSC), UAS-lacz(BDSC), UAS-BrpD3-mCherrypHlourin (gift from R. Hiesinger), UAS-myr-mCherrypHlourin (Jin *et al*., 2018), UAS-EGFR GFP (Zschätzsch *et al*., 2014), UAS-Brp B3.C8 RNAi (Knapek, Sigrist and Tanimoto, 2011), UAS-Syd1 RNAi(BDSC), UAS-Rab7 RNAi (VDRC), UAS-Atg6 RNAi (VDRC), UAS-Trans Tango (Talay *et al*., 2017), UAS-EGFR CA (Zschätzsch *et al*., 2014), UAS-tubulin Gal80^ts^(BDSC), Pacman Brp (Huang *et al*., 2020), UAS-Wasp RNAi(VDRC), UAS-Ensconsin RNAi(VDRC), UAS-Act42A RNAi(VDRC). Additional fly stocks used are w1118, valium 20, valium 10, canton-S as genetic controls (BDSC).

### Immunohistochemistry and Fixed Imaging

Pupal and adult brains were dissected in cold Schneider’s Drosophila medium and fixed in 4% paraformaldehyde (PFA) in PBS for ∼20 minutes. Tissues were then washed in PBST (1% Triton-X) on a shaker for 3×15mins followed by overnight incubation with primary antibodies at 4°C shaker. The primary antibodies used in this study were used with the following dilutions: chicken anti-GFP(1:250)(abcam), mouse anti-GFP (1:500)(Life Technology), goat anti-GFP (1:500)(abcam), goat anti-mCherry (1:500)(Sicgen), rabbit anti-Dsred (1:200)(Takara), mouse anti-Lacz (1:200)(Promega), rabbit anti-CD4 (1:250)(Invitrogen), rabbit anti-Rab7 (1:1000)(gift from Patrick Dolph). Next, the brains were washed again with 1% PBST on a shaker for 3×15mins. The brains were incubated with appropriate secondary antibodies (Alexa 488, 554, 647, 405, 594) at a concentration of 1:500 from Jackson ImmunoResearch Laboratories for 5-6hours at room temperature, followed by final wash with 1% PBST for 3×15mins. Tissues were mounted on taped cover slides using vector shield. Images were obtained with a Leica TCS SP8-X white laser confocal microscope with a 63x glycerol objective (NA=1.3)(Jin *et al*., 2018; Kiral *et al*., 2021).

### STED imaging

Adult brains were dissected in cold Schneider’s Drosophila medium and fixed in 4% paraformaldehyde (PFA) in PBS for ∼20 minutes. Tissues were then in PBST (1% Triton-X) on a shaker for 3×15mins followed by overnight incubation with primary antibodies at 4°C shaker. The primary antibodies used in this study with given dilutions were as follows: chicken anti-GFP (1:500) (abcam), mouse anti-Nc82 (1:10) (DSHB), rabbit anti-CD4(1:200) (Invitrogen). Next, the brains were washed again with 1% PBST on a shaker for 3×15mins. The brains were incubated in secondary antibodies: anti-ck STAR RED (1:250) (Abberior dyes), anti-mouse Alexa594 (1:100) (Thermofischer Scientific), anti-rabbit Alexa 488(1:100) (Thermofischer Scientific) for 5-6hours, followed by final wash with 1% PBST for 3×15mins. Then they were mounted on taped cover slides using Prolong Gold (Invitrogen) and kept 24hours at RT in dark. The slides were then stored at 4°C for 48hours before imaged. Images were obtained with a STED Expert Line Microscope from Abberior Instruments with a 100x oil objective (NA=1.4)(Pooryasin *et al*., 2021).

### Pupal brain culture and Live-Imaging

For all ex-vivo live imaging experiments, pupal or adult brain was carefully dissected out of the pupal case or the surrounding exoskeleton respectively. The resultant eye-brain complexes were mounted in 0.4% dialyzed low-melting agarose in a modified culture medium as described before (Özel *et al*., 2015). We used double sided tapes cut into 1inch x 1inch small squares as coverslips. Since all our developmental imaging were done after P48, we used Hydroxyecdysone free culture media. To fully expose DCN branch projection patterns, the pupae were mounted posterior side up. Live imaging was performed at room temperature using a Leica TCS SP8 X confocal microscope with a resonant scanner, using 63X water objective (NA=1.2), and optimized settings of minimal white laser excitation and crosstalk avoiding SP detector emission windows. White laser excitation was set to 488 nm for GFP, 554 nm for tdTomato signal acquisitions.

### Trans-tango and activity-dependent GRASP

Trans-tango was performed with DCN-specific ato-Gal4-14a (Hassan Bassem A., 2000)and M-DCN specific Gal4 (Linneweber *et al*., 2020)whereas GRASP experiment was performed with DCN-specific ato-LexA(Langen *et al*., 2013). Trans-tango flies were raised both at 18°C and 25°C to optimize the dissection conditions. 7 days old flies raised at 25°C showed dense connectivity pattern. The number of postsynaptic neurons was counted manually from their cell bodies using the “surface” tool in IMARIS, including all cell bodies with weak or strong labelling to reveal all potential connections. Since postsynaptic partner labeling by Trans-tango is age-dependent, 3-days old flies reared at 25°C were dissected for sparse labeling to reveal the identity of post-synaptic cell types connected to M-DCNs.

For activity-dependent GRASP experiments, to activate DCNs, freshly eclosed flies were transferred to 25°C incubator with 12-12hours light-dark cycle for 5 days. Brains were dissected and stained with a polyclonal anti-GFP antibody to label DCN pre-synaptic sites, monoclonal anti-GFP antibody to label GRASP signal, and polyclonal anti-CD4 antibody to label postsynaptic neurons(Kiral *et al*., 2021).

### Buridan’s paradigm assay

Fly navigation behavior was tested in a Buridan’s paradigm arena (Linneweber *et al*., 2020) using flies grown in a 12/12 hours light–dark cycle at 50% relative humidity. The arena consists of a round platform of 117 mm in diameter, surrounded by a water-filled moat and placed inside a uniformly illuminated white cylinder. The light was produced by four circular fluorescent tubes (Osram, L 40w, 640 C circular cool white) powered by an Osram Quicktronic QT-M 1 × 26–42. The fluorescent tubes were located outside of a diffuser (DeBanier, Belgium, 2090051, Kalk transparent, 180 g, white) positioned 147.5 mm away from the arena center. The temperature on the platform was kept constant at 25 °C. 30 mm-wide stripes of black cardboard were placed on opposing sides inside of the diffuser and served as visual targets. The retinal size of the visual object depends on the position of the fly on the platform. In this arena it ranges from 8.4° to 19.6° in width (11.7° in the center of the platform). Fly tracks were analyzed using CeTrAn (coulomb) and custom-written python code (Linneweber *et al*., 2020). 25 partially overlapping behavioral parameters were evaluated. Significant differences between experiment and controls and the rescue experiment were only found for parameters affecting motility.

The three selected activity related behavioral parameters are the following:

1. **Distance traveled (mm/min)**: Total distance travelled in mm per minute
2. **Activity time (s)**: Time active per minute in seconds
3. **Number of walks**: The number a fly walks from one stripe to the other. The fly needs to be on both ends near the edge closer than 80% of the platform radius for a walk to count.

### Quantification and statistical analysis

#### Branch number and length analysis

All imaging data were analyzed and presented with Imaris 9.0.1 (Bitplane). Branch numbers were detected automatically with the filament module using identical parameters for all experimental conditions (largest dendrite diameter: 3.0 µm, thinnest dendrite diameter: 0.2 µm). The resultant branch numbers were then recorded directly from the statistics tab of filament module and normalized it to the total number of axons per optic lobe. Any inconsistencies in automatic detection of branches were checked and corrected manually. Branch lengths were calculated manually using the “automatic placement” version of the filament module to calculate the 3D length of all branches. Intersection of axon shaft-primary branch were considered as the starting node and a filament was drawn till the respective branch tip. The resultant values of branch lengths were taken and recorded directly from the statistics tab of the filament module. Graph generation and statistical analyses were done using GraphPad Prism 8.2.0

#### Synapse number analysis

All imaging data were analyzed and presented with Imaris 9.0.1 (Bitplane). For synapse number analysis, CD4-tomato channel was used to generate surfaces for DCN axonal branches. Brp-positive puncta inside the surface were filtered using the masking function and were detected manually for individual branches. To obtain synapse distribution, we normalized the number of Brp-positive puncta inside individual DCN branch to the respective branch length which was calculated using the filament module as discussed above. Graph generation and statistical analyses were done using GraphPad Prism 8.2.0

#### Live tracing of molecules in branches

All imaging data were analyzed and presented with Imaris 9.0.1 (Bitplane) and the background noise was corrected with the threshold > background subtraction with a filter width of 60um in Imaris. GFP ‘+’ ve puncta were then tracked individually and manually for all the branches marked in CD4 channel over time and recorded. To obtain a distribution, we normalized the number of GFP-positive puncta inside individual DCN branch to the respective branch length which was calculated using the filament module. Graph generation and statistical analyses were done using GraphPad Prism 8.2.0

#### mCherry/pHluorin intensity analysis

Intensity analysis was performed using the surface module of Imaris 9.0.1 (Bitplane). All mCherry positive BrpD3 puncta were used to generate surface using the same threshold parameters (Diameter of largest spere which fits into the object=0.700 um; surface detail=0.481um) for experiments and controls. Mean intensity of the individual red channel (mCherry) to green channel (pHluorin) within each surface were recorded. Graph generation and statistical analyses were done using GraphPad Prism 8.2.0.

#### Colocalization analysis

All imaging data were analyzed and presented with Imaris 9.0.1 (Bitplane). For colocalization analysis, CD4-tomato channel was used to generate surfaces for DCN axonal branches. Brp-positive puncta (green channel) and the Rab7 puncta (red channel) inside the surface were filtered using the “masking” tool of surface module. All co-localization events were quantified manually on slice-by-slice basis for the entire z-stack in 2D. Only discernible individual compartments were counted. Partial or full correlation were given a score of 1 (“yes” colocalization) whereas 0 (“no” colocalization) if not. To obtain the fraction of colocalized events, the total number of colocalized Brp-Rab7 puncta were divided by the total BrpD3 puncta per axon. Graph generation and statistical analyses were done using GraphPad Prism 8.2.0.

### Statistical Analysis

Statistical comparison of two groups was performed with non-parametric Mann-Whitney test (T-test). Statistical comparison of more than two groups was performed with non-parametric Kruskal-Wallis test and corrected for multiple comparisons with Dunn’s as a post-hoc test. All significance values are denoted on the graphs and in their respective legends. Graph generation and statistical analyses were done using GraphPad Prism 8.2.0.

## Figure legends

**Supplementary 1.**
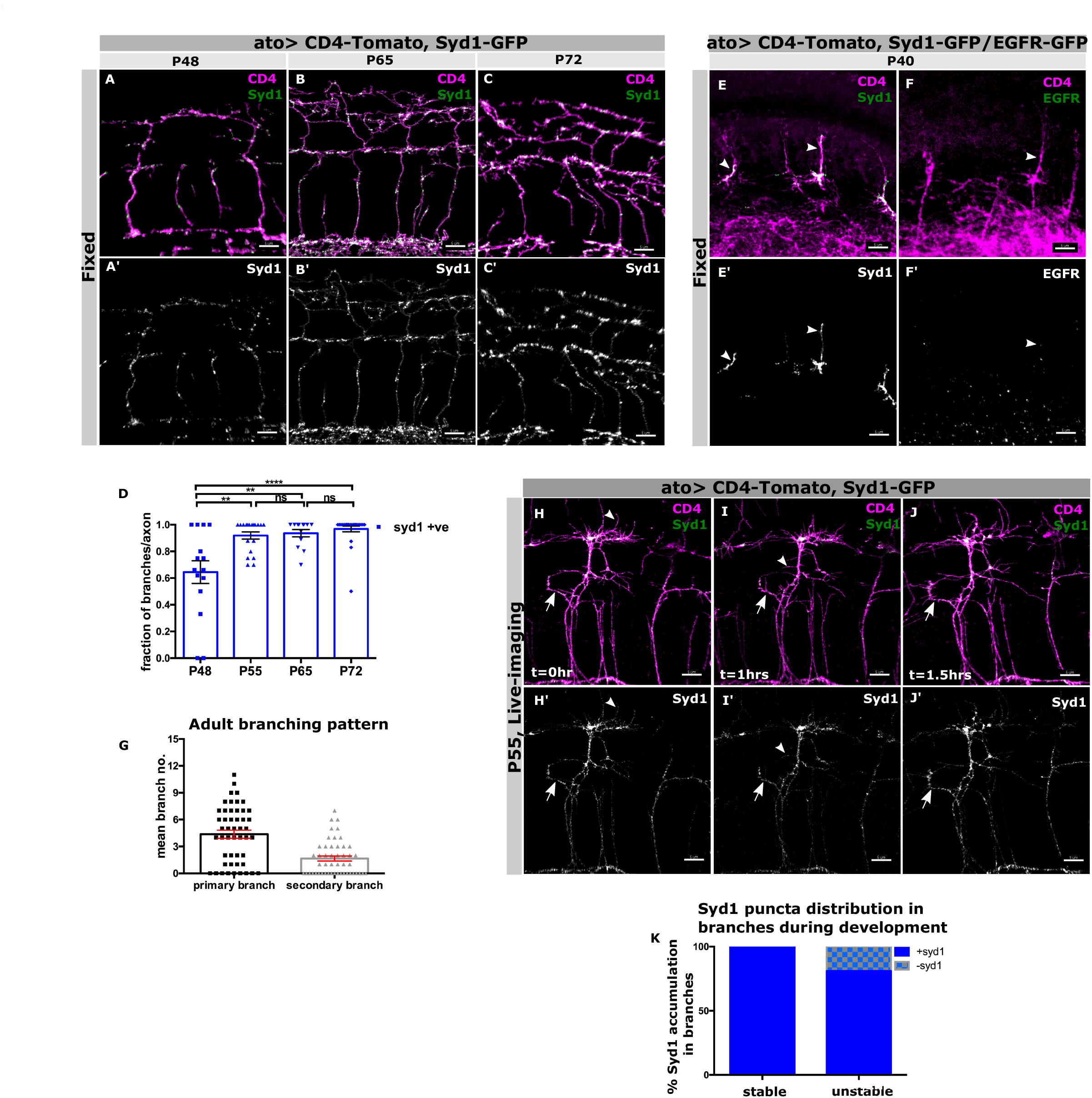
Spatio-temporal trafficking of Syd1 in the DCN branches during pupal development: **(A/A’-C/C’)** Temporal order of recruitment of Syd1 GFP (green) in the DCN branches (magenta) during development; P48 **(A/A’)**, P65 **(B/B’)** and P72 **(C/C’). (D)** Quantification showing the fraction of DCN branches containing Syd1 puncta per axon during development. N=15 axons at P48, N=19 axons at P55, N=13 axons at P65 and N=25 axons at P72; \*\*\*\**p* < 0.0001, Mann Whitney test. **(E/E’-F/F’)** Syd1 GFP (green)(arrowhead) puncta **(E/E’)** gets trafficked prior to EGFR GFP (green)(arrowhead)**(F/F’)** in DCN axonal shafts (magenta) at P40. **(G/G’-I/I’)** Spatial distribution pattern of Syd1 GFP (green) in DCN branches (magenta) during development in ex-vivo pupal brain cultures between P55-P57. Z-stack projection of ex-vivo time-lapse imaging of control axons showing Syd1 GFP (green) localize in both stable (arrow) and unstable branches (arrowhead). **(J)** Quantification showing percentage of stable (blue) and unstable (checkered blue) branches containing Syd1 puncta during development. N=25 stable branches and N=36 unstable branches. Error bars denote mean±SEM, scalebar represents 5μm.

**Supplementary Video 1**

**Live trafficking of Syd1, EGFR and BrpD3 in the DCN branches during pupal development:**

Live-imaging of EGFR GFP, BrpD3 GFP and Syd1 GFP in DCN branches labeled by membrane marker CD4 Tomato driven by atoGal4-14a starting P55.

**Supplementary 2.**
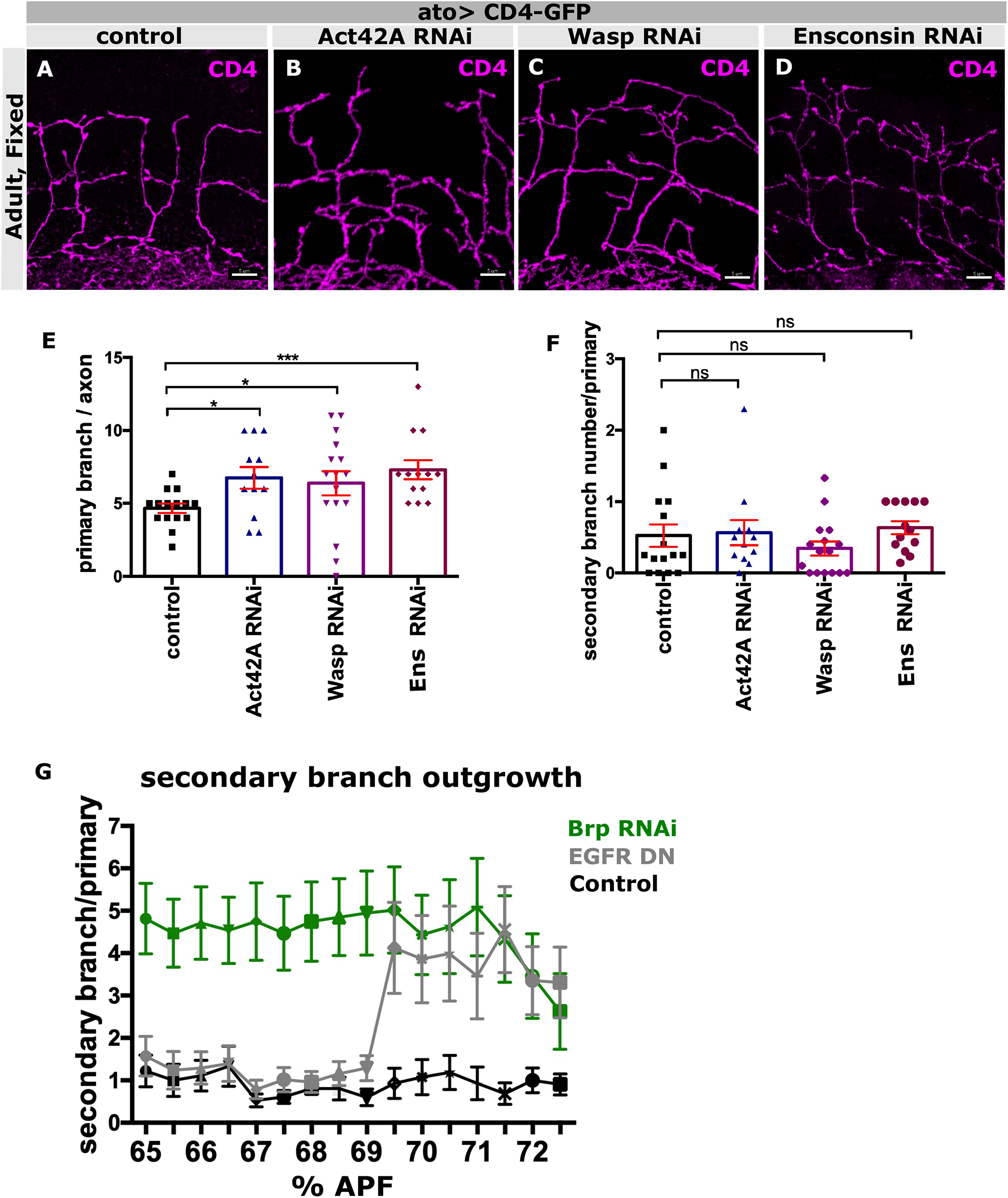
Effect of knocking down cytoskeletal proteins on axon branching; temporal outgrowth of secondary branches: **(A-D)** Adult DCN branch morphology (magenta) upon knocking down several cytoskeletal regulators like Act42A **(B)**, Wasp **(C)**, Ensconsin(Ens) **(D**) compared to genetic control **(A).(E)** Adult quantification showing increased number of primary branches per axon in uas-Act42A RNAi (blue), uas-Wasp RNAi (pink), uas-Ens RNAi (purple) compared to control (black). N=15 axons for control, N=12 axons for Act42A RNAi, N=16 axons for Wasp RNAi and N=13 axons for Ens RNAi; \**p=*0.0231, \**p=*0.0294, \*\*\**p=*0.0003, Mann Whitney test. **(F)** Adult quantification showing unaffected secondary branches normalized to primary branch number in uas-Act42A RNAi (blue), uas-Wasp RNAi (pink), uas-Ens RNAi (purple) compared to control (black). N=15 axons for control, N=12 axons for Act42A RNAi, N=16 axons for Wasp RNAi and N=13 axons for Ens RNAi; ^**ns**^*p=*0.5059, ^**ns**^*p=*0.6402, ^**ns**^*p=*0.1545, Mann Whitney test. **(G)** Quantification of the temporal outgrowth of secondary branches (secondary branch number normalized to primary branches) in every 30mins interval between P65-P72 ex-vivo cultures in Brp B3,C8 RNAi (green) or EGFR DN (gray) compared to control (black). N=29 branches for Brp B3,C8 RNAi (green); N=20 branches for EGFR DN (gray) and N=18 branches for control (black). ^ns^*p* = 0.6867 for control, ^ns^*p* = 0.9441 for Brp B3,C8 RNAi and \*\*\*\**p* < 0.0001 for EGFR DN. Error bars denote mean±SEM, scalebar represents 5μm.

**Supplementary Video 2.1**

**Ext./retr. dynamics of adult DCN branches in control vs Brp KD:**

Live-imaging of adult (^∼^5-7 days old) DCN branches labeled by atoGal4-14a driving membrane marker uas-CD4 GFP in control vs Brp B3,C8 RNAi.

**Supplementary Video 2.2**

**Secondary branch outgrowth in DCN branches in control vs Brp KD and EGFR DN:**

Live-imaging of DCN branches labeled by atoGal4-14a driven membrane marker uas-CD4 GFP in control vs Brp B3,C8 RNAi and EGFR DN post P65.

**Supplementary 3.**
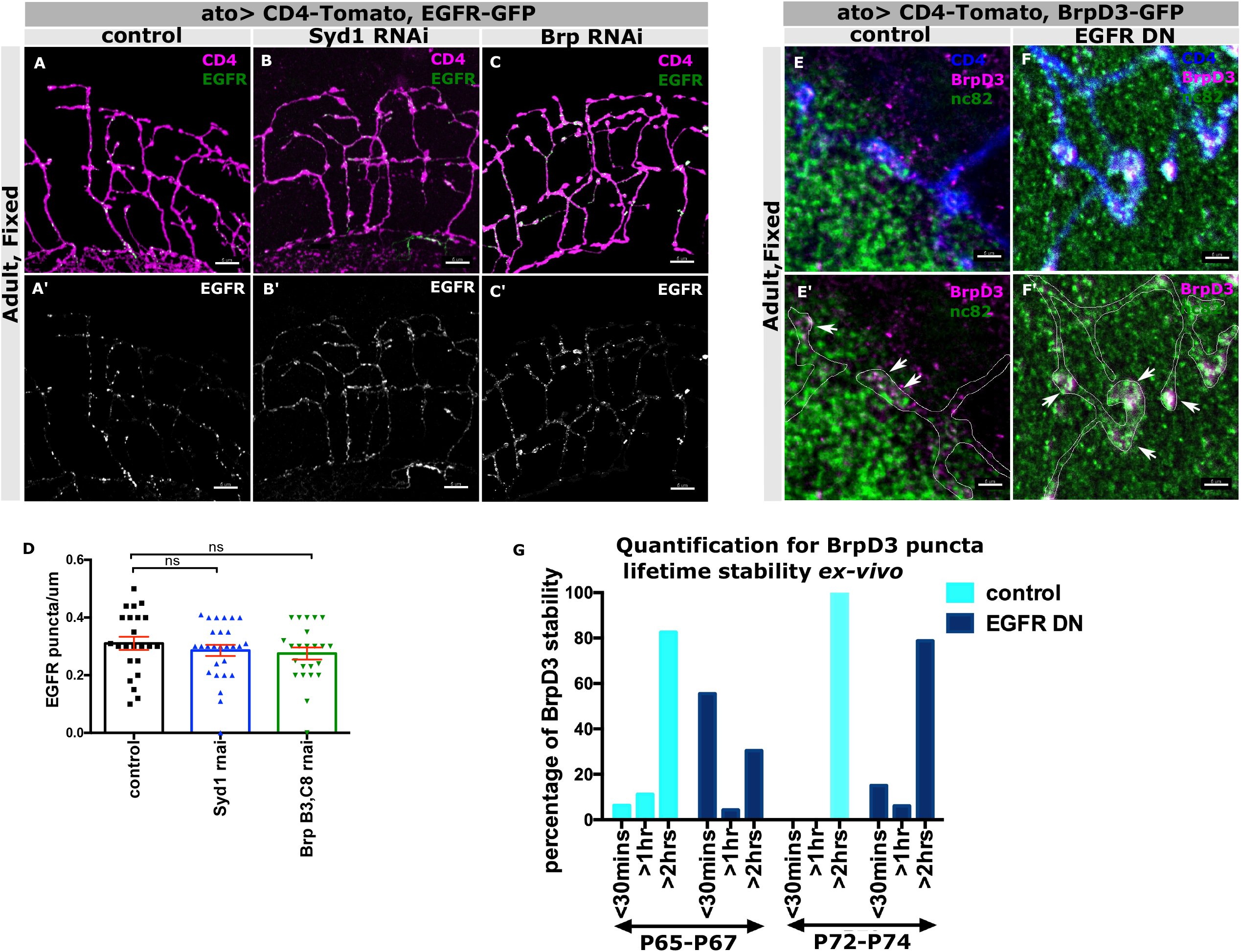
Genetic hierarchy of EGFR, Syd1 and BrpD3; co-localization of endogenous Brp with uas-BrpD3 GFP; temporal stability of BrpD3 GFP punctum in control vs EGFR DN. **(A/A’-C/C’)** EGFR-GFP (green) localization in adult DCN branches (magenta) upon knocking down Syd1 **(B)**, Brp **(C)** compared to control **(A). (D)** Quantification showing unaffected distribution of EGFR GFP puncta per unit branch length in Syd1 RNAi (blue), Brp B3,C8 RNAi (green) compared to control (black) in adults. N=23 branches for control, N=27 branches for Syd1 RNAi and N=23 branches for Brp RNAi; ^**ns**^*p=*0.4714, ^**ns**^*p=*0.2612. Mann Whitney test. **(E/E’-F/F’)** STED co-localization analysis of endogenous Brp marked by Nc82 staining (green) and BrpD3 GFP (magenta) in DCN axon branches (red) in control **(E/E’)** vs EGFR DN **(F/F’). (G)** Quantification of percentage of BrpD3-GFP puncta lifetime (<30mins, <1hour, >2hours) in individual DCN branches tracked for 2hours from P65 and P72 in control (black) vs EGFR DN (gray). N=20 punctum for control and 23 punctum, for EGFR DN in P65-P67 ex-vivo culture. N=28 punctum for control and 33 punctum for EGFR DN in P72-P74 ex-vivo culture. Error bars denote mean±SEM, scalebar represents 5μm.

**Supplementary 4.**
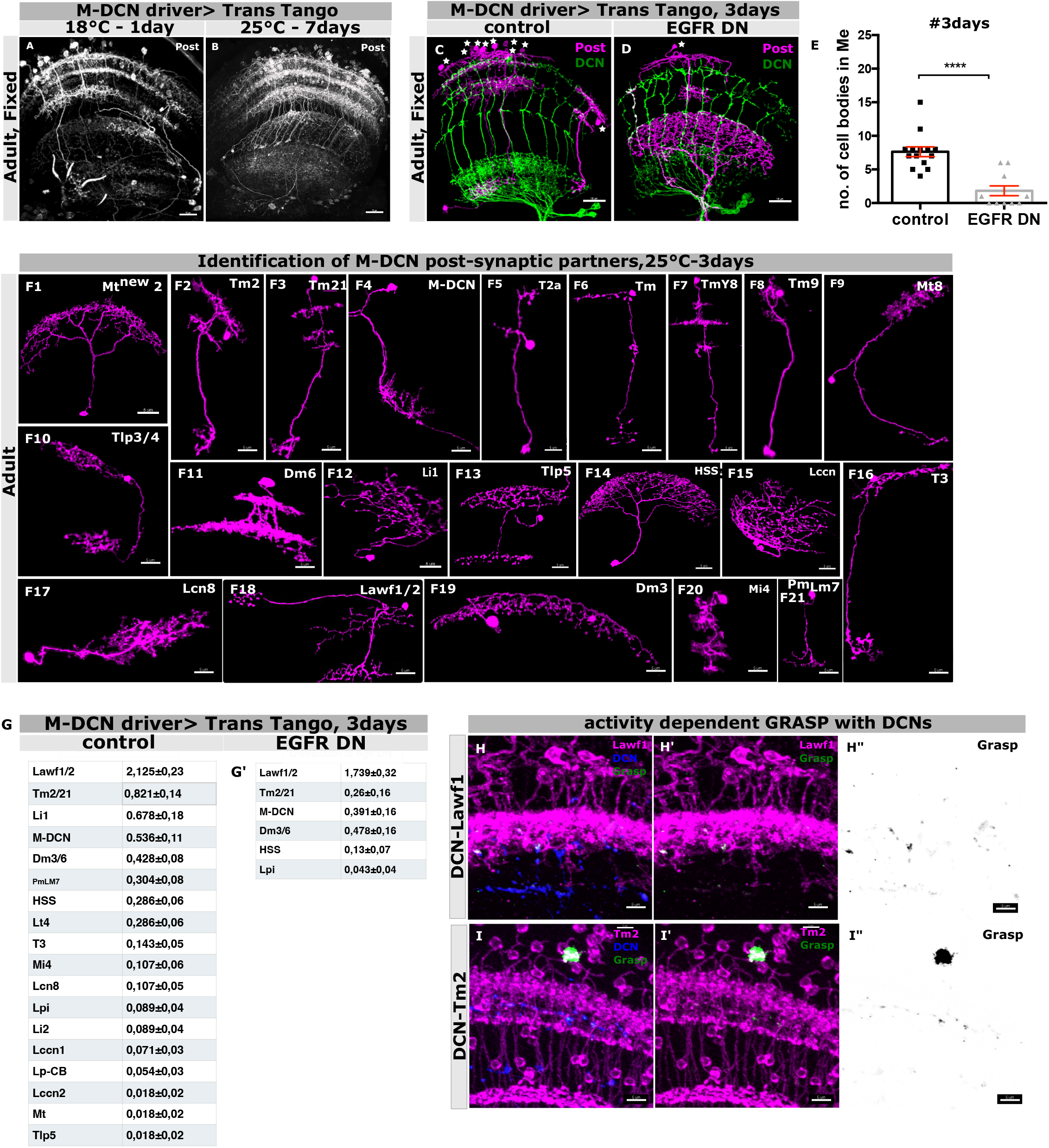
Loss of EGFR activity causes a reduction in M-DCN circuit connectivity: **(A-B)** Optimization for labeling of post-synaptic partners of adult M-DCNs driven by a M-DCN specific Gal4 using Trans-Tango at 18°C for 1day old adults **(A)** and 25°C for 7days old adults **(B). (C-D)** Sparse labeling of post-synaptic partners (magenta) of adult M-DCNs (green) driven by a M-DCN specific Gal4 using Trans-Tango at 25°C for 3 days old adults in control **(C)** vs EGFR DN **(D). (E)** Quantification showing reduced number of post-synaptic cell bodies in medulla per optic lobe in control (black) vs EGFR DN (gray) in adult DCNs at 25°C for 3 days. N=14 optic lobes for control, N=11 optic lobes for EGFR DN. Kruskal–Wallis and Dunn’s as post-hoc test; \*\*\*\**p* < 0.0001. **(F1-F21)** Identification of post-synaptic partners (magenta) of adult DCNs using Trans-Tango at 25°C for 3 days using a M-DCN specific neuronal driver. We characterized and identified several post-synaptic targets like Mt^new^2 **(F1)**, Tm2 **(F2)**, Tm21**(F3)**, M-DCN **(F4)**, T2a **(F5)**, Tm**(F6)**, TmY8 **(F7)**, Tm9 **(F8)**, Mt8 **(F9)**, Tlp3/4 **(F10)**, Dm6 **(F11)**, Li1 **(F12)**, Tlp5 **(F13)**, HSS **(F14)**, Lccn **(F15)**, T3 **(F16)**, Lcn8 **(F17)**, Lawf1/2**(F18)**, Dm3 **(F29)**, Mi4 **(F20)**, Pm_LM7_ **(F21)**. Table of Post-synaptic partner distribution of M-DCN in control **(G)** vs EGFR DN **(G’). (H/H”-I/I”)** Grasp signal (green) confirming synaptic connection between adult DCN pre-synaptic sites (blue) with Lawf1 (magenta) **(H-H”)** and Tm2 (magenta) **(I-I”)**. Ato-LexA is used to drive adult DCNs for this experiment. N= 50 optic lobes for control, N= 50 optic lobes for EGFR DN. Error bars denote mean±SEM, scalebar represents 20μm in **(A-B)**, 15 μm in **(C-D)** and 5μm in **(F1-21, H/H”-I/I”)** except **(F18)** which represents 10μm.

**Supplementary Video 3**

**Temporal stability of BrpD3 GFP punctum in control vs EGFR DN:**

Live-imaging of atoGal4-14a driven BrpD3 GFP in DCN branches marked by uas-CD4 Tomato starting P65 and P72 of pupal development in control vs EGFR DN.

**Supplementary 5.**
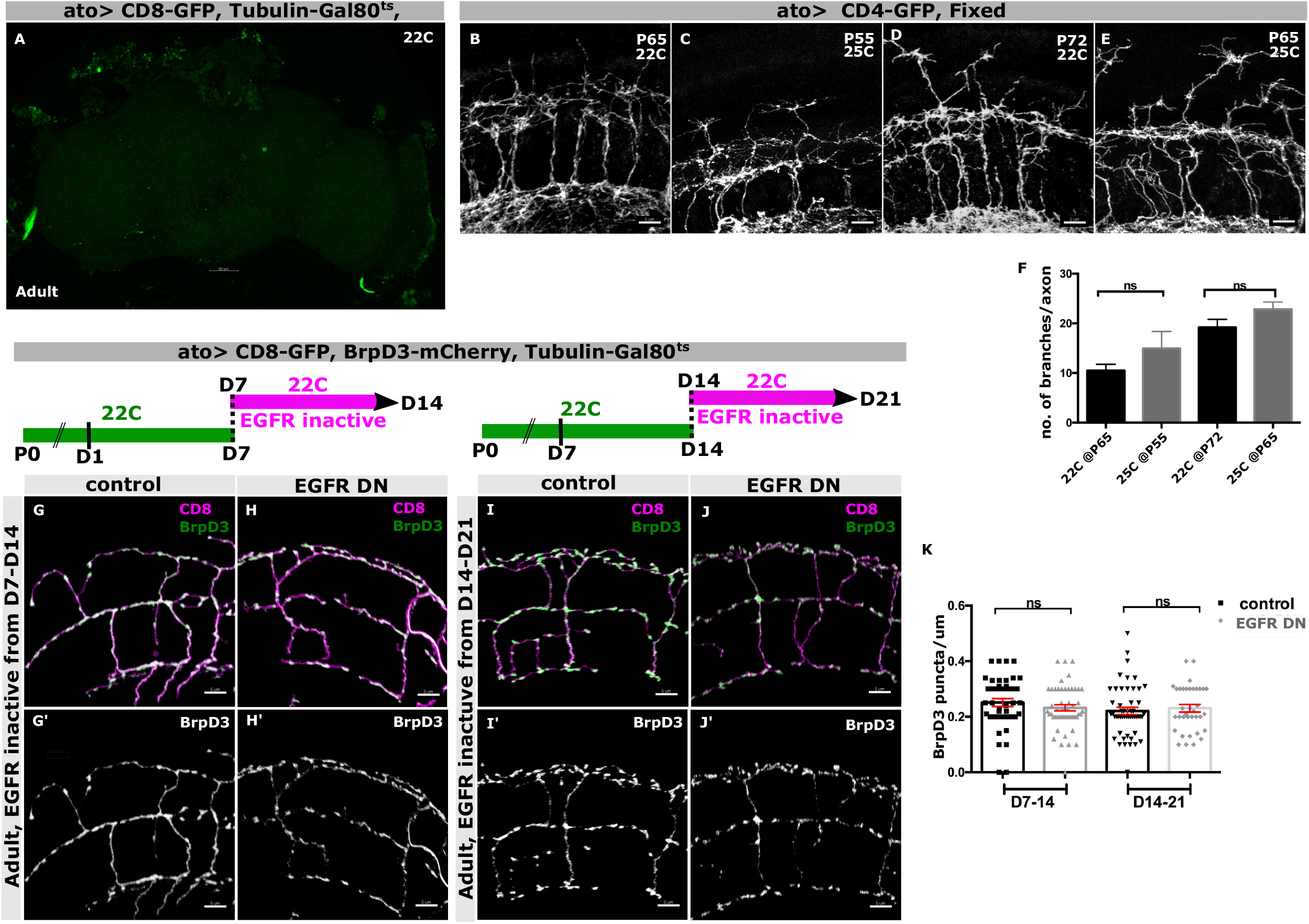
Experimental set up control and DCN developmental calibration of 22°C vs 25°C; adult specific requirement of EGFR activity for proper branch consolidation and synaptogenesis: **(A)** Control experiment for checking Gal80^ts^ efficiency in repressing *atoGal4-14a* driven uas-CD8 GFP expression when the fly was raised at 22°C throughout development. **(B-E)** Calibration of DCN axon branch progression at 22°C vs 25°C during development. In-vivo progression of DCN branch morphology (white) at 65hours APF at 22°C **(B)** vs 55hours APF at 25°C **(C)**, and 72hours APF at 22°C **(D)** vs 65hours APF at 25°C **(E). (F)** Quantification showing branch number per axon of 65hours APF and 72hours APF at 22°C (black) corresponds to 55hours APF and 65hours APF at 25°C (gray) respectively. N=5 lobes for P65 at 22°C, N=7 lobes for P55 at 25°C, N=5 lobes for P72 at 22°C and N=7 lobes for P65 at 22°C; ^**ns**^*p* =0.4476, ^**ns**^*p* =0.2476, Mann Whitney test. **(G/G’-J/J’)** Inactivating EGFR activity in adult DCN branches (green) only between D7-D14 **(G/G’-H/H’)** or between D14-D21 **(I/I’-J/J’)** did not affect BrpD3-mCherry (red) puncta localization in the branches. **(K)** Quantification showing unaltered number of BrpD3 GFP puncta per unit branch length in control (black) vs EGFR DN (gray) expressed during D7-D17 and D14-D21. N=44 branches for control D7-14, N=51 branches for EGFR DN D7-14, N=51 branches for control D14-21 and N=36 branches for EGFR DN D14-21; ^**ns**^*p* =0.1344 and ^**ns**^*p* =0.5053, Mann Whitney test. Error bars denote mean± SEM, scalebar represents 5μm whereas in **(A)** represents 50μm.

**Supplementary 6.**
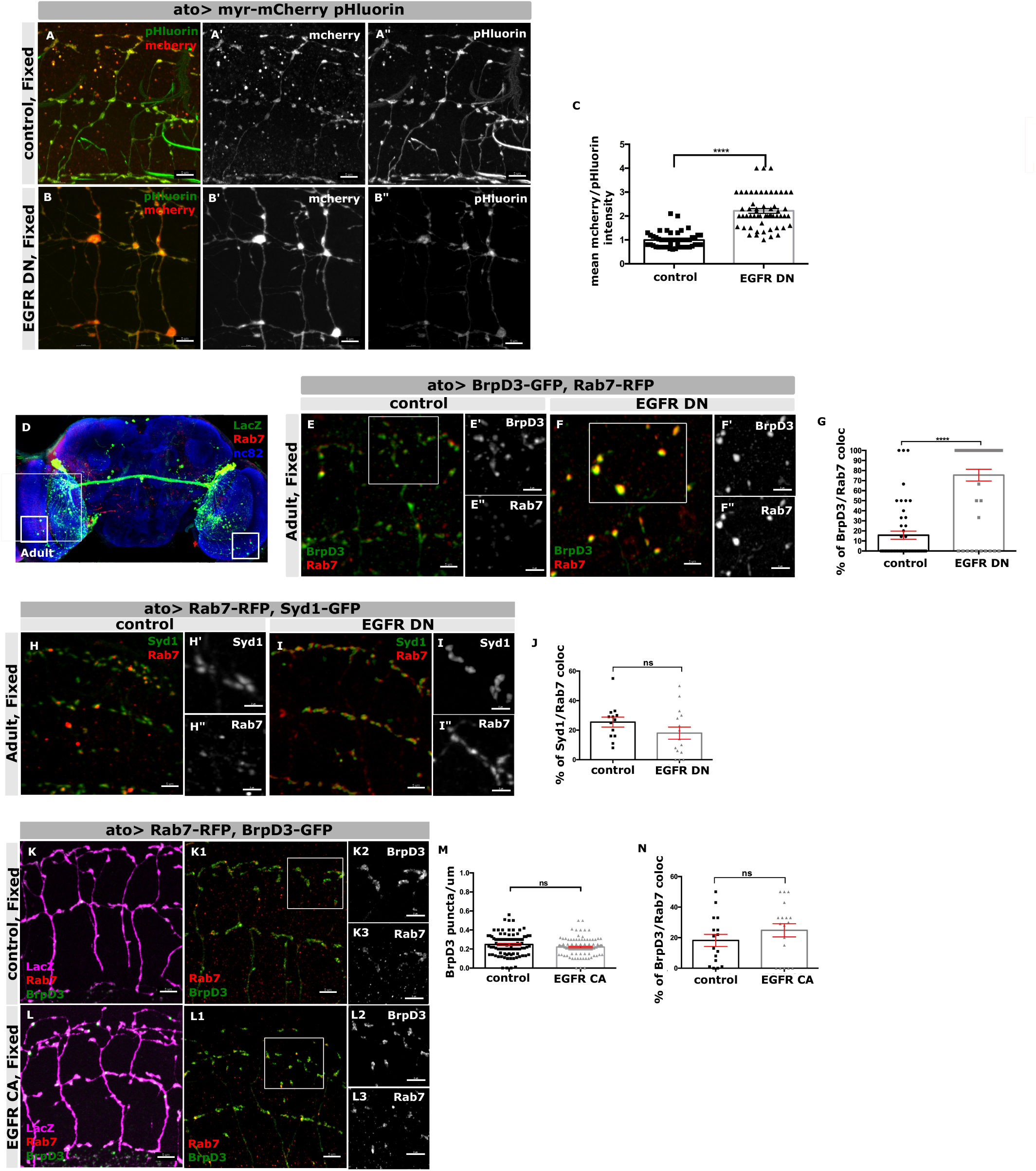
Lack of EGFR activity in adult DCNs causes degradation of general plasma membrane cargoes; Adult co-localization of Active Zones (AZ) with uas-Rab7 RFP in lack of EGFR activity; Syd1 GFP does not colocalize with late endosomes in lack of EGFR activity; DCNs expressing EGFR CA shows no AZs loss in branches with no increased co-localization with late endosomes: **(A/A”-B/B”)** Increase in mCherry signal (red) intensity compared to pHluorin (green) intensity when expressing the general membrane degradation probe uas-myr-mCherry-pHluorin in adult DCN branches in EGFR DN **(B-B”)** compared to control **(A-A”). (C)** Quantification showing increased mean intensity of mCherrry channel to pHluorin channel across several myr-DF regions per axon in adults. N=50 axons in control and 56 axons in EGFR DN. Mann Whitney test; \*\*\*\**p* < 0.0001. **(D)** Representative adult fly brain with neuropils marked by Nc82 (blue), DCNs labeled by LacZ (green) and late endosomes marked by Rab7 RFP (red) localized in the branches in medulla (white box). **(E/E”-F/F”)** Adult localization of BrpD3 GFP (green) and Rab7 RFP (red) in control **(E-E”)** vs EGFR DN **(F-F”)** in DCN branches. White box represents region shown in higher magnification **(E’/E”-F/F”). (G)** Quantification showing increased percentage of BrpD3 GFP puncta colocalized with Rab7 RFP puncta in the adult branches per axon in control (black) vs EGFR DN (gray). N=49 punctum for control, N=49 punctum for EGFR DN. Mann Whitney test; \*\*\*\**p* < 0.0001. **(H/H”-I/I”)** Adult localization of Syd1 GFP (green) and Rab7 RFP in control **(H-H”)** vs EGFR DN **(I-I”)** in DCN branches. **(J)** Adult quantification showing unaffected percentage of Syd1 GFP puncta colocalized with Rab7 RFP puncta in DCN branches per axon in control (black) vs EGFR DN (gray). N=13 axons for control and N=15 branches for EGFR DN; ^**ns**^*p=*0.1386, Mann Whitney test. **(K/K3-L/L3)** Adult DCN branches (magenta) exhibits BrpD3 GFP (green) and Rab7 RFP (red) puncta in control **(K)** vs EGFR CA **(L). (M)** Adult quantification showing unaffected number of BrpD3 GFP puncta per unit branch in control (black) vs EGFR CA (gray). N=90 branches for control and N=114 branches for EGFR CA; ^**ns**^*p*=0.0590, Mann Whitney test. **(N)** Quantification showing unaltered percentage of BrpD3 GFP puncta colocalized with Rab7 RFP puncta per axon in control (black) vs EGFR CA (gray) adults. Individual dots represent single axons. N=16 branches in control and N=18 branches in EGFR CA; ^**ns**^*p* =0.3750, Mann Whitney test. Error bars denote mean± SEM, scalebar in **(D)**represents 50μm, **(A-A”, B-B”, E, F, H, I, K, L)** represents 5μm and **(E’/E’’, F’/F’’, H’/H’’, I’/I’’**,**K2/K3, L2/L3)** represents 3μm.

**Supplementary 7.**
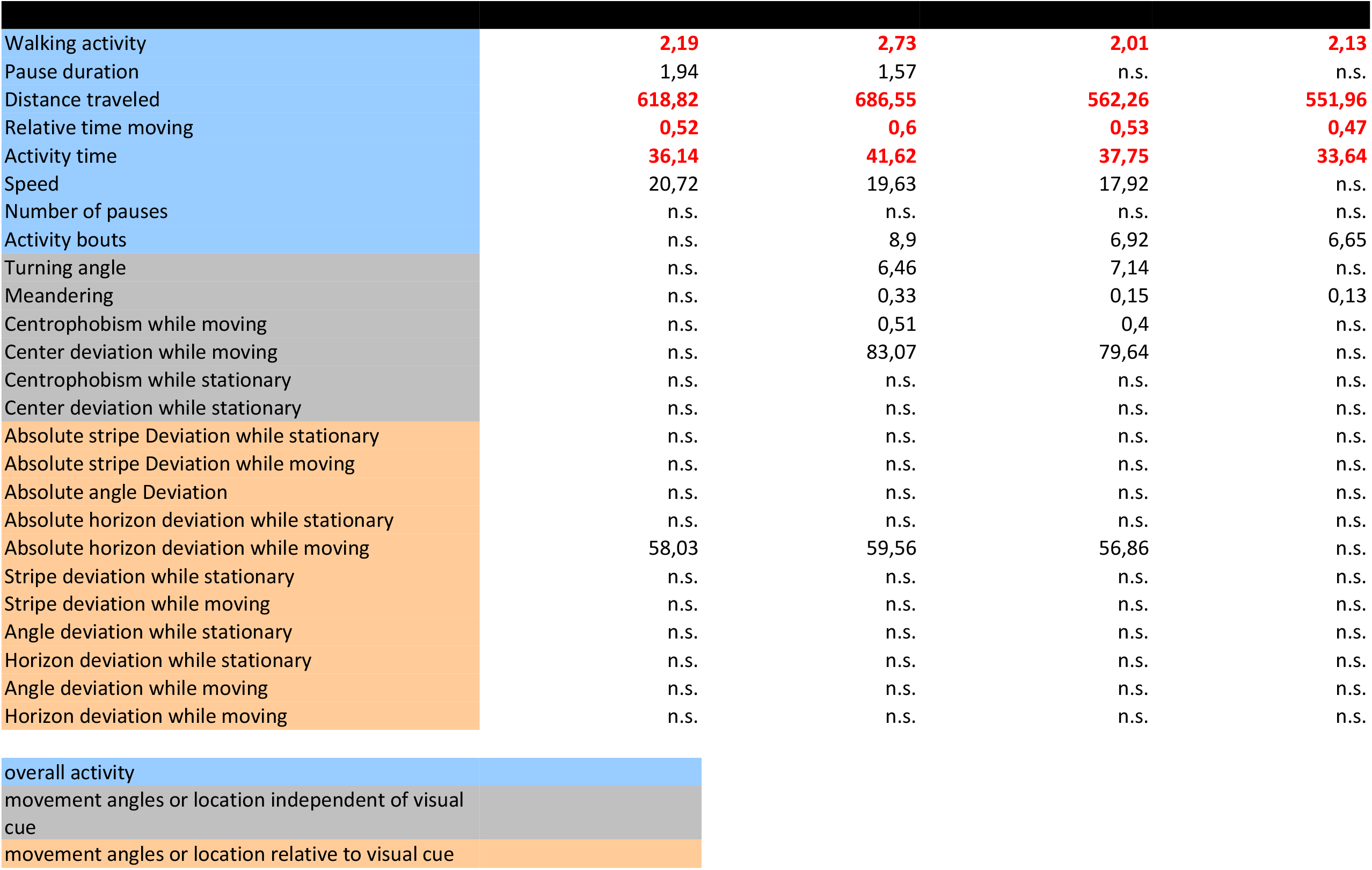
Table of parameters measured for individual flies in Buridan’s paradigm.

## References

Adnan, G. et al. (2020) ‘The GTPase Arl8B plays a principle role in the positioning of interstitial axon branches by spatially controlling autophagosome and lysosome location’, Journal of Neuroscience, 40(42), pp. 8103–8118. doi: 10.1523/JNEUROSCI.1759-19.2020.

Agi, E., Kulkarni, A. and Hiesinger, P. R. (2020) ‘Neuronal strategies for meeting the right partner during brain wiring’, Current Opinion in Neurobiology. Elsevier Ltd, pp. 1–8. doi: 10.1016/j.conb.2020.01.002.

Aguirre, A., Rubio, M. E. and Gallo, V. (2010) ‘Notch and EGFR pathway interaction regulates neural stem cell number and self-renewal’, Nature, 467(7313). doi: 10.1038/nature09347.

Azarnia Tehran, D., Kuijpers, M. and Haucke, V. (2018) ‘Presynaptic endocytic factors in autophagy and neurodegeneration’, Current Opinion in Neurobiology. doi: 10.1016/j.conb.2017.12.018.

Batool, S. et al. (2019) ‘Synapse formation: From cellular and molecular mechanisms to neurodevelopmental and neurodegenerative disorders’, Journal of Neurophysiology, 121(4). doi: 10.1152/jn.00833.2018.

Bhukel, A. et al. (2019) ‘Autophagy within the mushroom body protects from synapse aging in a non-cell autonomous manner’, Nature Communications, 10(1). doi: 10.1038/s41467-019-09262-2.

Bourgeois, J. P. and Rakic, P. (1993) ‘Changes of synaptic density in the primary visual cortex of the macaque monkey from fetal to adult stage’, Journal of Neuroscience, 13(7). doi: 10.1523/jneurosci.13-07-02801.1993.

Buff, E. et al. (1998) ‘Signalling by the Drosophila epidermal growth factor receptor is required for the specification and diversification of embryonic muscle progenitors’, Development, 125, pp. 2075–2086. Available at: https://cob.silverchair-cdn.com/cob/content_public/journal/dev/125/11/10.1242_dev.125.11.2075/1/2075.pdf?Expires=1643013366&Signature=Awz9f59jhGNfKXWb6R∼a8TKyq4bTBpNcpY9fGb0gLNkmi2SOb0wJJeIRDS6btlZ3IYH5AhSBe98nZguOHyAhY-1uI9q2waRfU0qfbiQxb7pCJFqdIaYz0SVxL.

Chen, B. E. et al. (2006) ‘The Molecular Diversity of Dscam Is Functionally Required for Neuronal Wiring Specificity in Drosophila’, Cell, 125(3). doi: 10.1016/j.cell.2006.03.034.

Chia, P. H. et al. (2014) ‘Local F-actin network links synapse formation and axon branching’, Cell, 156(1–2). doi: 10.1016/j.cell.2013.12.009.

Cisneros-Franco, J. M. et al. (2020) ‘Critical periods of brain development’, in Handbook of Clinical Neurology. Elsevier B.V., pp. 75–88. doi: 10.1016/B978-0-444-64150-2.00009-

Colón-Ramos, D. A. (2009) ‘Chapter 2 Synapse Formation in Developing Neural Circuits’, Current Topics in Developmental Biology. doi: 10.1016/S0070-2153(09)01202-2.

Constance, W. D. et al. (2018) ‘Neurexin and neuroligin-based adhesion complexes drive axonal arborisation growth independent of synaptic activity’, eLife, 7. doi: 10.7554/eLife.31659.

Doll, C. A. and Broadie, K. (2014) ‘Impaired activity-dependent neural circuit assembly and refinement in autism spectrum disorder genetic models’, Frontiers in Cellular Neuroscience, 8(FEB). doi: 10.3389/fncel.2014.00030.

Economo, M. N. et al. (2016) ‘A platform for brain-wide imaging and reconstruction of individual neurons’, eLife, 5(JANUARY2016). doi: 10.7554/eLife.10566.

Feller, M. B. and Scanziani, M. (2005) ‘A precritical period for plasticity in visual cortex’, Current Opinion in Neurobiology, 15(1), pp. 94–100. doi: 10.1016/J.CONB.2005.01.012.

Fischbach, K. F. and Dittrich, A. P. M. (1989) ‘The optic lobe of Drosophila melanogaster. I. A Golgi analysis of wild-type structure’, Cell and Tissue Research, 258(3), pp. 441–475. doi: 10.1007/BF00218858.

Fleming, A. and Rubinsztein, D. C. (2020) ‘Autophagy in Neuronal Development and Plasticity’, Trends in Neurosciences. doi: 10.1016/j.tins.2020.07.003.

Fouquet, W. et al. (2009) ‘Maturation of active zone assembly by Drosophila Bruchpilot’, Journal of Cell Biology, 186(1), pp. 129–145. doi: 10.1083/jcb.200812150.

Galvez-Contreras, A. Y., Quiñones-Hinojosa, A. and Gonzalez-Perez, O. (2013) ‘The role of EGFR and ErbB family related proteins in the oligodendrocyte specification in germinal niches of the adult mammalian brain’, Frontiers in Cellular Neuroscience, 7(DEC). doi: 10.3389/fncel.2013.00258.

Goldshmit, Y. et al. (2004) ‘SOCS2 Induces Neurite Outgrowth by Regulation of Epidermal Growth Factor Receptor Activation’, Journal of Biological Chemistry, 279(16). doi: 10.1074/jbc.M312873200.

Grueber, W. B. et al. (2005) ‘The development of neuronal morphology in insects’, Current Biology. doi: 10.1016/j.cub.2005.08.023.

Hassan, B. A. and Hiesinger, P. R. (2015) ‘Beyond Molecular Codes: Simple Rules to Wire Complex Brains’, Cell, 163(2). doi: 10.1016/j.cell.2015.09.031.

Hassan Bassem A. (2000) atonal Regulates Neurite Arborizationbut Does Not Act as a Proneural Genein the Drosophila Brain.

Hiesinger, P. R. (2021) ‘Brain wiring with composite instructions’, BioEssays, 43(1). doi: 10.1002/BIES.202000166.

Hiesinger, P. R. and Hassan, B. A. (2018) ‘The Evolution of Variability and Robustness in Neural Development’, Trends in Neurosciences, 41(9), pp. 577–586. doi: 10.1016/j.tins.2018.05.007.

Hoersting, A. K. and Schmucker, D. (2021) ‘Axonal branch patterning and neuronal shape diversity: roles in developmental circuit assembly: Axonal branch patterning and neuronal shape diversity in developmental circuit assembly’, Current Opinion in Neurobiology. Elsevier Ltd, pp. 158–165. doi: 10.1016/j.conb.2020.10.019.

Hua, J. Y. and Smith, S. J. (2004) ‘Neural activity and the dynamics of central nervous system development’, Nature Neuroscience. doi: 10.1038/nn1218.

Huang, S. et al. (2020) ‘Presynaptic Active Zone Plasticity Encodes Sleep Need in Drosophila’, Current Biology, 30(6), pp. 1077–1091.e5. doi: 10.1016/j.cub.2020.01.019.

Huber, K. M. et al. (2015) ‘Dysregulation of mammalian target of rapamycin signaling in mouse models of autism’, Journal of Neuroscience, 35(41), p. 13836. doi: 10.1523/JNEUROSCI.2656-15.2015.

Jin, E. J. et al. (2018) ‘Live Observation of Two Parallel Membrane Degradation Pathways at Axon Terminals’, Current Biology, 28(7), pp. 1027–1038.e4. doi: 10.1016/j.cub.2018.02.032.

Kalil, K. and Dent, E. W. (2014) ‘Branch management: Mechanisms of axon branching in the developing vertebrate CNS’, Nature Reviews Neuroscience. doi: 10.1038/nrn3650.

Kim, M. et al. (2016) ‘Mutation in ATG5 reduces autophagy and leads to ataxia with developmental delay’, eLife, 5(JANUARY 2016). doi: 10.7554/eLife.12245.001.

Kiral, F. R. et al. (2020) ‘Autophagy-dependent filopodial kinetics restrict synaptic partner choice during Drosophila brain wiring’, Nature Communications, 11(1). doi: 10.1038/s41467-020-14781-4.

Kiral, F. R. et al. (2021) ‘Brain connectivity inversely scales with developmental temperature in Drosophila’, Cell Reports, 37(12), p. 110145. doi: 10.1016/j.celrep.2021.110145.

Knapek, S., Sigrist, S. and Tanimoto, H. (2011) ‘Bruchpilot, A synaptic active zone protein for anesthesia-resistant memory’, Journal of Neuroscience, 31(9), pp. 3453–3458. doi: 10.1523/JNEUROSCI.2585-10.2011.

Konstantinides, N. et al. (2018) ‘Phenotypic Convergence: Distinct Transcription Factors Regulate Common Terminal Features’, Cell, 174(3), pp. 622–635.e13. doi: 10.1016/j.cell.2018.05.021.

Koprivica, V. et al. (2005) ‘Neuroscience: EGFR activation mediates inhibition of axon regeneration by myelin and chondroitin sulfate proteoglycans’, Science, 310(5745). doi: 10.1126/science.1115462.

Langen, M. et al. (2013) ‘Mutual inhibition among postmitotic neurons regulates robustness of brain wiring in Drosophila’, eLife, 2013(2). doi: 10.7554/eLife.00337.

Lewis, T. L. et al. (2018) ‘MFF-dependent mitochondrial fission regulates presynaptic release and axon branching by limiting axonal mitochondria size’, Nature Communications, 9(1). doi: 10.1038/s41467-018-07416-2.

Linneweber, G. A. et al. (2020) ‘A neurodevelopmental origin of behavioral individuality in the Drosophila visual system’, Science, 367(6482). doi: 10.1126/science.aaz4547.

Menzies, F. M. et al. (2017) ‘Autophagy and Neurodegeneration: Pathogenic Mechanisms and Therapeutic Opportunities’, Neuron. doi: 10.1016/j.neuron.2017.01.022.

Meyer, M. P. and Smith, S. J. (2006) ‘Evidence from in vivo imaging that synaptogenesis guides the growth and branching of axonal arbors by two distinct mechanisms’, Journal of Neuroscience, 26(13). doi: 10.1523/JNEUROSCI.0223-06.2006.

Moyer, C. E., Shelton, M. A. and Sweet, R. A. (2015) ‘Dendritic spine alterations in schizophrenia’, Neuroscience Letters. doi: 10.1016/j.neulet.2014.11.042.

Niell, C. M. (2006) ‘Theoretical analysis of a synaptotropic dendrite growth mechanism’, Journal of Theoretical Biology, 241(1), pp. 39–48. doi: 10.1016/j.jtbi.2005.11.014.

Owald, D. et al. (2010) ‘A Syd-1 homologue regulates pre-and postsynaptic maturation in Drosophila’, Journal of Cell Biology, 188(4), pp. 565–579. doi: 10.1083/jcb.200908055.

Özel, M. N. et al. (2015) ‘Filopodial dynamics and growth cone stabilization in Drosophila visual circuit development’, eLife, 4(OCTOBER2015). doi: 10.7554/eLife.10721.

Özel, M. Neset et al. (2019) ‘Serial Synapse Formation through Filopodial Competition for Synaptic Seeding Factors’, Developmental Cell, 50(4), pp. 447–461.e8. doi: 10.1016/j.devcel.2019.06.014.

Özel, M Neset et al. (2019) ‘Serial Synapse Formation through Filopodial Competition for Synaptic Seeding Factors’, Developmental Cell, 50(4), pp. 447–461.e8. doi: 10.1016/j.devcel.2019.06.014.

Pooryasin, A. et al. (2021) ‘Unc13A and Unc13B contribute to the decoding of distinct sensory information in Drosophila’, Nature Communications, 12(1). doi: 10.1038/s41467-021-22180-6.

Poultney, C. S. et al. (2013) ‘Identification of small exonic CNV from whole-exome sequence data and application to autism spectrum disorder’, American Journal of Human Genetics, 93(4), pp. 607–619. doi: 10.1016/j.ajhg.2013.09.001.

Powchik, P. et al. (1998) ‘Postmortem studies in schizophrenia’, Schizophrenia Bulletin, 24(3). doi: 10.1093/oxfordjournals.schbul.a033330.

Rajgor, D., Welle, T. M. and Smith, K. R. (2021) ‘The Coordination of Local Translation, Membranous Organelle Trafficking, and Synaptic Plasticity in Neurons’, Frontiers in Cell and Developmental Biology. Frontiers Media S.A. doi: 10.3389/fcell.2021.711446.

Rico, B. et al. (2004) ‘Control of axonal branching and synapse formation by focal adhesion kinase’, Nature Neuroscience, 7(10). doi: 10.1038/nn1317.

Ruthazer, E. S., Li, J. and Cline, H. T. (2006) ‘Stabilization of axon branch dynamics by synaptic maturation’, Journal of Neuroscience, 26(13). doi: 10.1523/JNEUROSCI.0069-06.2006.

Schmid, A. et al. (2008) ‘Activity-dependent site-specific changes of glutamate receptor composition in vivo’, Nature Neuroscience, 11(6), pp. 659–666. doi: 10.1038/nn.2122.

Sênos Demarco, R. and Jones, D. L. (2020) ‘EGFR signaling promotes basal autophagy for lipid homeostasis and somatic stem cell maintenance in the Drosophila testis’, Autophagy, 16(6), pp. 1145–1147. doi: 10.1080/15548627.2020.1739450.

Shao, Z. et al. (2019) ‘Dysregulated protocadherin-pathway activity as an intrinsic defect in induced pluripotent stem cell–derived cortical interneurons from subjects with schizophrenia’, Nature Neuroscience, 22(2), pp. 229–242. doi: 10.1038/s41593-018-0313-z.

Shen, W. and Ganetzky, B. (2009) ‘Autophagy promotes synapse development in Drosophila’, Journal of Cell Biology, 187(1). doi: 10.1083/jcb.200907109.

Simmons, A. B. et al. (2017) ‘DSCAM-mediated control of dendritic and axonal arbor outgrowth enforces tiling and inhibits synaptic plasticity’, Proceedings of the National Academy of Sciences of the United States of America, 114(47). doi: 10.1073/pnas.1713548114.

Spinner, M. A., Walla, D. A. and Herman, T. G. (2018) ‘Drosophila syd-1 has rhogap activity that is required for presynaptic clustering of bruchpilot/elks but not neurexin-1’, Genetics, 208(2), pp. 705–716. doi: 10.1534/genetics.117.300538.

Srahna, M. et al. (2006) ‘A signaling network for patterning of neuronal connectivity in the Drosophila brain’, PLoS Biology, 4(11). doi: 10.1371/journal.pbio.0040348.

Stavoe, A. K. H. and Holzbaur, E. L. F. (2019) ‘Autophagy in neurons’, Annual Review of Cell and Developmental Biology. doi: 10.1146/annurev-cellbio-100818-125242.

Tagliatti, E. et al. (2016) ‘Arf6 regulates the cycling and the readily releasable pool of synaptic vesicles at hippocampal synapse’, eLife, 5. doi: 10.7554/eLife.10116.

Talay, M. et al. (2017) ‘Transsynaptic Mapping of Second-Order Taste Neurons in Flies by trans-Tango’, Neuron, 96(4), pp. 783–795.e4. doi: 10.1016/j.neuron.2017.10.011.

Tang, G. et al. (2014) ‘Loss of mTOR-Dependent Macroautophagy Causes Autistic-like Synaptic Pruning Deficits’, Neuron, 83(5), pp. 1131–1143. doi: 10.1016/j.neuron.2014.07.040.

Ting, C. Y. et al. (2011) ‘Focusing transgene expression in drosophila by Coupling Gal4 with a Novel Split-Lex A Expression System’, Genetics, pp. 229–233. doi: 10.1534/genetics.110.126193.

Urwyler, O. et al. (2019) ‘Branch-restricted localization of phosphatase Prl-1 specifies axonal synaptogenesis domains’, Science, 364(6439). doi: 10.1126/science.aau9952.

Vaughn, J. E., Barber, R. P. and Sims, T. J. (1988) ‘Dendritic development and preferential growth into synaptogenic fields: A quantitative study of Golgi-impregnated spinal motor neurons’, Synapse, 2(1). doi: 10.1002/syn.890020110.

Vijayan, V. and Verstreken, P. (2017) ‘Autophagy in the presynaptic compartment in health and disease’, Journal of Cell Biology. doi: 10.1083/jcb.201611113.

Wagh, D. A. et al. (2006) ‘Bruchpilot, a protein with homology to ELKS/CAST, is required for structural integrity and function of synaptic active zones in Drosophila’, Neuron, 49(6), pp. 833–844. doi: 10.1016/j.neuron.2006.02.008.

Wang, M. M. et al. (2019) ‘The relationship between autophagy and brain plasticity in neurological diseases’, Frontiers in Cellular Neuroscience, 13. doi: 10.3389/fncel.2019.00228.

Winkle, C. C. et al. (2016) ‘Beyond the cytoskeleton: The emerging role of organelles and membrane remodeling in the regulation of axon collateral branches’, Developmental Neurobiology. doi: 10.1002/dneu.22398.

Wu, M. and Zhang, P. (2020) ‘EGFR-mediated autophagy in tumourigenesis and therapeutic resistance’, Cancer Letters. Elsevier Ireland Ltd, pp. 207–216. doi: 10.1016/j.canlet.2019.10.030.

Xiong, W. et al. (2020) ‘Autophagy is Required for Remodeling in Postnatal Developing Ribbon Synapses of Cochlear Inner Hair Cells’, Neuroscience, 431, pp. 1–16. doi: 10.1016/J.NEUROSCIENCE.2020.01.032.

Xu, Y. and Quinn, C. C. (2016) ‘Transition between synaptic branch formation and synaptogenesis is regulated by the lin-4 microRNA’, Developmental Biology, 420(1), pp. 60–66. doi: 10.1016/j.ydbio.2016.10.010.

Zschätzsch, M. et al. (2014) ‘Regulation of branching dynamics by axon-intrinsic asymmetries in Tyrosine Kinase Receptor signaling’, eLife, 3. doi: 10.7554/elife.01699.

